# Evolution of genetic variance and its consequences for eco-evolutionary responses in complex mutualistic networks

**DOI:** 10.1101/2024.12.25.630074

**Authors:** Gaurav Baruah, Meike J Wittmann

**Author notes:** These authors contributed equally.

## Abstract

Rising temperatures threaten biodiversity and ecosystem resilience by disrupting phenological synchrony in plant–pollinator networks. These interactions, essential for ecosystem functioning, are sensitive to temperature shifts. We model species eco-evolutionary responses to an abrupt temperature increase, comparing evolution in mean trait alone versus evolution involving both the mean and genetic variance. We show that networks where species evolve in both their mean phenotype and genetic variance exhibit increased resilience to temperature shifts. Hyper-generalist species, engaged in extensive mutualistic interactions, accumulate greater genetic variance, thereby facilitating faster evolutionary rescue under strong selective pressures. Specialists also benefit by interacting with these hyper-generalists, which stabilize adaptation to new temperatures. We observe emergence in opposing selection pressures in complex networks that promote increases in genetic variance (“evolvability”), reducing trait lag, enabling faster adaptation, and increasing survival. Our findings highlight the role of evolving genetic variance and network architecture in mitigating plant–pollinator phenotypic mismatches under rising environmental temperatures. This study provides insights into the adaptive capacity of mutualistic networks, highlighting dynamic genetic variance changes as key to resilience amid accelerating climate warming. Our study highlights the importance of feedbacks between network topology and genetic architecture for conservation under climate change.

## 1 Introduction

The increasing challenges posed by climate change, particularly rising temperatures, have been shown to have negative impacts on the life cycles of organisms (Baruah *et al.*, 2021; Hegland *et al.*, 2009; Opedal, 2019), leading to alterations in ecological communities. As temperatures increase, complex communities must adapt not only to the changing thermal environment (Scranton & Amarasekare, 2017) but also to shifts in the life cycles of interaction partners embedded within these complex communities (Baruah & Lakämper, 2024; Freimuth *et al.*, 2022; Hegland *et al.*, 2009). The rate of these adaptations will inevitably influence the resilience of such ecological systems (Baruah & Lakämper, 2024). This is true for predator-prey relationships (Yamamichi & Miner, 2015), aquatic ecosystems (Bolchoun *et al.*, 2017; Morgan *et al.*, 2020) and, critically, plant-pollinator systems (Baruah & Lakämper, 2024; Baruah & Wittmann, 2024; Duchenne *et al.*, 2020; Hegland *et al.*, 2009). Plant-pollinator networks among others are especially vulnerable to changes in environmental temperatures, as abrupt temperature increases can disrupt the synchrony of phenological events (Baruah & Lakämper, 2024; Duchenne et al., 2020; Freimuth et al., 2022; Hegland et al., 2009; Labonté et al., 2023; Trunschke et al., 2024). Species may be able to survive these events through evolutionary adaptation to the altered environment (Bell & Gonzalez, 2009; Klausmeier et al., 2020; Opedal, 2019). However, species within complex communities face selective pressures to adapt not only to the new environment but also to changes in interactions with other species in the community. Investigating adaptation to altered environments without considering complex species interactions will provide an incomplete understanding of the process.

Plant-pollinator interactions are integral to the reproductive success of many plants and are key to maintaining biodiversity (Bascompte & Scheffer, 2023; Bascompte & Stouffer, 2009; Hegland *et al.*, 2009; Labonté et al., 2023; Opedal, 2019). As temperatures rise, mismatches between plants and their pollinators can result in decreased pollination success and, consequently, diminished plant reproduction (Labonté *et al.*, 2023). This vulnerability is particularly pronounced in insect-pollinated plants, as their emergence is intricately tied to pollinators whose life cycles are directly influenced by temperature (Hegland *et al.*, 2009; Menzel *et al.*, 2006; Miller-Rushing & Primack, 2008). Furthermore, climate-induced shifts in plant-pollinator phenological events can push these vital species to the brink of decline or weaken species interactions that can cause critical transition to undesirable states such as a collapse of the entire network (Baruah & Lakämper, 2024). The vulnerability of plant-pollinator communities to collapse has broader implications for biodiversity, ecosystem stability, and the provision of essential ecosystem services.

Eco-evolutionary dynamics play a pivotal role in shaping the complex relationships between species, driving their co-evolution as well as adaptation to environmental changes. One important factor that plays an important role in maintaining complex relationship in plantpollinator networks is variation within species (Arroyo-Correa *et al.*, 2023; Baruah, 2022; Baruah & Wittmann, 2024; Cantwell-Jones et al., 2024). In the context of plant-pollinator interactions, there is intraspecific variation in traits such as phenology, where individuals exhibit variation in their emergence or flowering onset (Hegland *et al.*, 2009). Multiple processes contribute to this variation, including a genetic component (Opedal, 2019). As anthropogenic changes drive increases in temperatures, understanding the eco-evolutionary changes in traits impacting plant-pollinator interactions becomes paramount. The variation in these traits can significantly influence species interactions within mutualistic communities. However, there is currently no theoretical framework or understanding of how such trait variation could evolve in response to multiple selective pressures, such as shifts in the environment, changes in competitive pressure from species within a guild, or alterations in mutualistic partners, all of which can change simultaneously when local environmental temperature, i.e. the temperate experienced by the community, shifts. Hypothetically, a strong directional shift in environmental conditions could reduce genetic variance within species, potentially hindering adaptation to new conditions. However, this scenario becomes more complex in the context of multiple species interactions within a large plant-pollinator network. It is also possible that due to the presence of various species interactions (both positive and negative), a significant shift in environmental temperature could potentially also increase trait variance in species, also commonly known as “evolvability” (Houle, 1992; Opedal, 2019). This might occur due to conflicting selective pressures: one driving species to remain matched with their mutualistic partners, and the other pushing them to adapt to the new environmental conditions while also competing with others. These dynamics could ultimately shape the resilience of complex plant-pollinator networks in novel environments. However, these hypotheses lack theoretical support at present.

The complex relationship between the structure and function of plant-pollinator networks requires an additional focus on the consequences of the structural aspects of these interactions on species persistence. By adopting a network theory approach, one can define the connectance of a specific plant-pollinator network as the proportion of the number of established connections between mutualistic partners in comparison to the total number of potential interactions between the guilds of plants and pollinators. Network connectance is a characteristic tied to the architecture of plant-pollinator communities, and it is crucial to consider it in addition with the total number of species. Another crucial concept inherent to plant-pollinator networks is their nestedness. In the context of plant-pollinator networks, nestedness means that a limited number of hyper-generalist species exist, whose set of interaction partners encompasses the set of partners of more specialized species (Bascompte *et al.*, 2003; Bascompte & Stouffer, 2009). Such nestedness is often evident across multiple levels in plant pollinator networks (Guimarães *et al.*, 2017). Beyond mitigating the risk of extinction, the high number of interactions involving a generalist species appears to confer a selective advantage on their mutualistic partners, including specialists (Baruah, 2022; Bascompte *et al.*, 2003). The positive impact of having a greater number of mutualistic partners extends to the whole of the plant-pollinator network. Previous studies have demonstrated that the architecture of a plant-pollinator network as quantified by connectance or nestedness appears to enhance the resilience of plant-pollinator networks (Baruah, 2022; Baruah & Lakämper, 2024; Bastolla et al., 2009). Given all these considerations, the architecture of such plant-pollinator networks is bound to interact with eco-evolutionary dynamics and thereby impact species responses to temperature increases.

In this study, we develop a theoretical framework to assess how complex mutualistic plantpollinator networks can undergo evolutionary adaptation through genetic evolution, despite intricate relationships with both partner species and competitors. In such mutualistic plantpollinator interactions, phenology and variation in phenology, such as insect emergence and/or plant flowering, play a critical role in successful plant-pollinator interactions. We present a theoretical model in which we model such a trait that is dependent on environmental temperature. In addition, we model how the variation in such a trait can evolve under strong selection pressures due to a shift in the local environmental temperature, and examine how such a shift might influence eco-evolutionary adaptation of species.

We introduce a quantitative eco-evolutionary framework based on modified generalized Lotka-Volterra equations, allowing for the evolution of the trait mean and its genetic variance. We then compare scenarios where 1) only the mean trait evolves, 2) both mean trait and genetic variance evolve, and 3) species do not evolve at all, to understand their relative impacts on adaptation to novel environmental temperatures. We disentangle the influence of eco-evolutionary factors and network architecture on species survival, showing that trait variance, competitive pressure, and a species’ network position - defined by its degree and proximity to a super-generalist— shape network resilience.

## 2 Methods and Materials

### 2.1 Modelling framework: Plant-pollinator mutualistic dynamics

We model the dynamics of pollinators 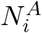 and plants 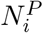 in an ecologically relevant quantitative trait *z* that relates to its optimum phenotype, i.e. the temperature that would be optimal for the individual. Here the superscripts *A* and *P* stand for pollinators and plants, respectively. Within each plant or pollinator species *i*, the trait is normally distributed as given by the probability density function 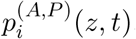, with mean *u*_*i*_ and phenotypic variance *ρ*_*i*_, with each individual having its own trait value *z*. The governing dynamical equations of population dynamics can be written with modified Lotka-Volterra equations. The per-capita growth rate of pollinators, 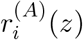, and plants, 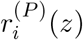, can be written as modified from Baruah & Lakämper (2024); Baruah & Wittmann (2024) *(see details in supplementary appendix):*

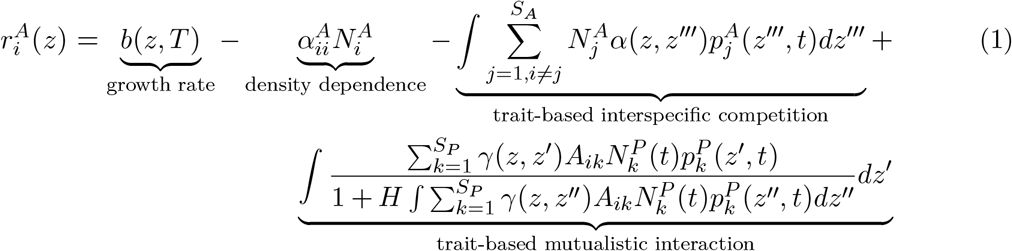

where, 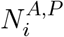 gives the density of species *i* belonging to either animals or plants. In this model, *b*(*z, T*) denotes the temperature-dependent growth rate of a specific phenotype, independent of competitive or mutualistic advantages for both animals and plants, each exhibiting a similar dependency. This implies that individuals within a guild will experience the impact of local temperature, denoted as *T*. A phenotype’s growth is also dependent on local environmental temperature, given by *b*(*z, T*). *S*_*A*_ and *S*_*P*_ are the number of pollinator and plant species, respectively. *H* represents the handling time, while *A*_*ik*_ signifies the corresponding entry in the adjacency matrix A, with a value of 1 indicating interaction between species *i* and *k*, and 0 representing no interaction. 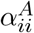 in equation 1 refers to density dependence of species *i* (here pollinator), and *α*(*z, z^′′′^*) refers to a Gaussian pairwise competition kernel between phenotypes of different species belonging to the same guild of species, i.e., either pollinators or plants. We assume that if different pollinators have similar phenotypes, for instance, they become the most active or emerge at similar temperatures, the intensity of interspecies competition for resources will increase. Consequently, the formulation becomes:

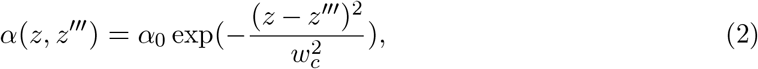

where *α*_0_ is the maximum interspecific competition intensity and *w*_*c*_ is the width of the interspecific competition kernel.

In addition, *γ*(*z, z^′^*) characterizes mutualistic interactions among individuals, where *z* denotes the optimum phenotype of a pollinator individual and *z*^*′*^ denotes the optimum phenotype of a plant individual. This function can also be modeled as a Gaussian function:

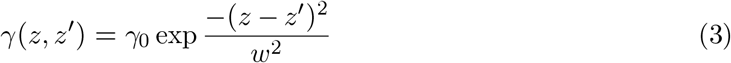

In this equation, *γ*_0_ represents the maximum strength of mutualistic interactions, while the width of the interaction kernel *w* controls the extent of interaction among individuals of different guilds of species. When the optimal phenotypes of a plant and a pollinator are similar, mutualistic benefits are enhanced. Biologically this could mean that the optimum flowering temperature of plants matches the optimum emergence temperature of pollinators, increasing the likelihood of mutualistic interaction (Baruah & Lakämper, 2024; Guimarães *et al.*, 2011; Thompson *et al.*, 2002). Temperature strongly influences species phenology, such as plant flowering times and pollinator emergence, which are often tied to monthly temperatures. Thus, mutualistic interactions are more likely when the optimum traits align (Hegland *et al.*, 2009).

The temperature-dependent growth or tolerance curve is derived from a modified version of the studies by Amarasekare & Johnson (2017); Baruah & Lakämper (2024):

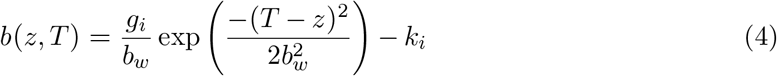

In this equation, *T* represents the local temperature, while *b*_*w*_ determines the width of the curve, and the ratio 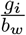 influences the height of the peak of the curve (Amarasekare & Johnson, 2017; Åkesson *et al.*, 2021). *k*_*i*_ is the extrinsic mortality constant. The trait of a species *i* thus is under selective pressure from changes in the temperature *T* and from the strength in inter-species competition and mutualistic interactions.

Taking everything together, the population dynamics of animal species *i* can be written as by integrating equation 1 over the trait space (see appendix 1):

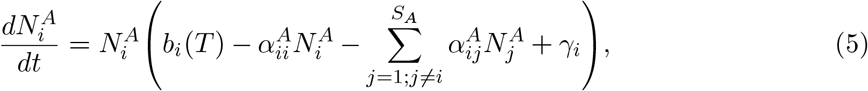

where 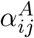 is the ecological impact of competition by pollinator species *j* on species *i* based on their mean optimum trait values, and *γ*_*i*_ is similarly the mutualistic impact of all plant species on pollinator *i*. Here, the exact formulation of 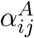 and *γ*_*i*_ is given in the supplementary section 1. Similarly, an analogous equation can be formulated for the plant species with index *A* replaced by *P*. The evolutionary dynamics of the mean optimum trait, *u*_*i*_, are given as:

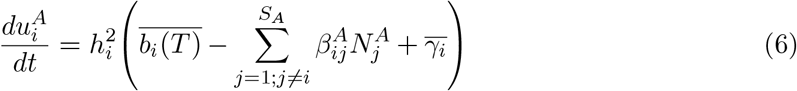

and evolutionary dynamics of genetic variance of the optimum phenotype,

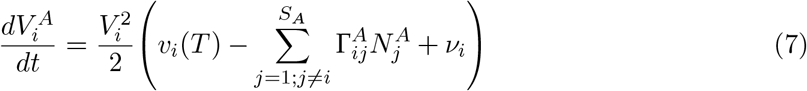

where *V*_*i*_ is the genetic variance of the optimum trait for species *i* in a mutualistic community. Phenotypic variance in our modelling setup was given as *ρ*_*i*_ = *V*_*i*_ + *ϵ*, where *ϵ* is the environmental variance. Thus, the broad sense heritability, 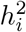, also changes as genotypic variance evolves in response to changes in interspecific competition, mutualistic interaction, and tolerance to temperature. Here, 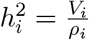 for a species *i* in question, 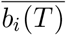 is the evolutionary impact of temperature on changes in the mean optimum trait 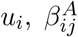 is the evolutionary impact of competition on change in mean optimum trait of species 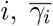 is the evolutionary impact of mutualistic trait-based interaction on changes in mean trait, *v*_*i*_(*T*), Γ_*ij*_, and ν_*i*_ are the effects of environmental temperature, competition and trait-based mutualistic interaction on changes in the genetic variance of species *i*, respectively.

## 3 Plant-pollinator networks and eco-evolutionary dynamics

To capture rapid declines in the fitness of species within a plant-pollinator network as a consequence of a shift in the environmental temperature, we modeled the environmental temperature change as a press perturbation, representing a permanent shift in the environment, *T*. A qualitative overview of the expected modelling outcomes is presented in Figure 1.

**Figure 1.**
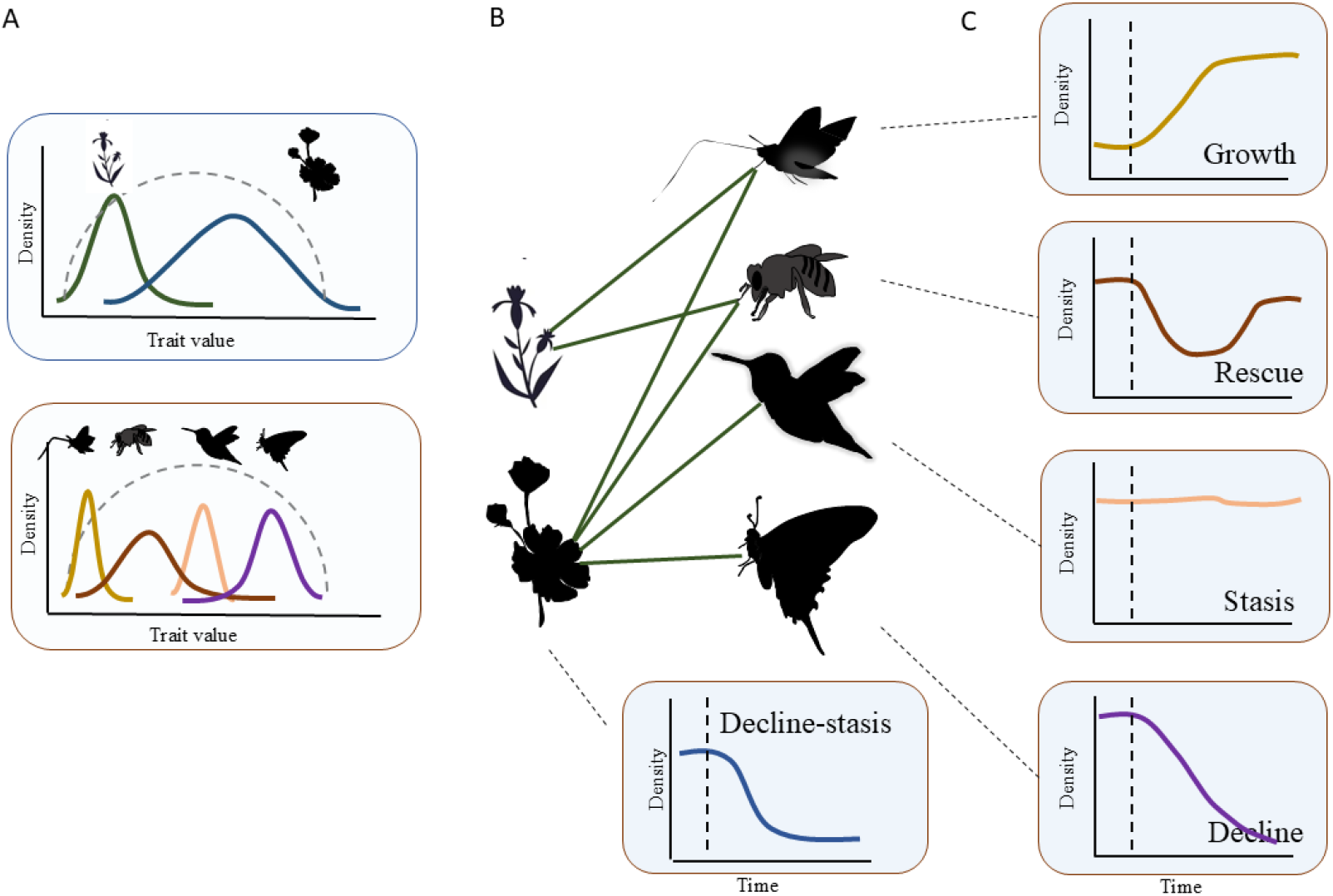
Example graphic of our model setup for a six-species plant–pollinator community (two plants, four pollinators) responding to an abrupt temperature shift. (A) Trait distributions at equilibrium before the shift, with dashed lines indicating region of positive growth. (B) Network structure showing variation in species’ interaction structure. (C) After the temperature shift (dashed vertical line), species can exhibit one of five dynamics: (1) Growth, (2) Rescue (initial decline followed by recovery), (3) Stasis, (4) Decline to extinction, or (5) Decline-stasis (drop followed by low stable density). We assess how these outcomes vary under three models: no evolution, evolution of mean trait only, and evolution of both mean trait and genetic variance.

To begin, we assembled 61 empirical plant-pollinator networks and their corresponding adjacency matrices, **A** (quantifying species interactions), sourced from the web-of-life database (www.web-of-life.es). These networks varied in size, containing between 8 and 70 species.

After extracting these plant-pollinator adjacency matrices, we simulated the evolution of these networks under a constant environmental temperature, (*T*), set at 22 °*C*, over a period of (2 * 103) time points. The initial conditions for the simulation included assigning random mean trait values to both plant and pollinator species, (*u*_*i*_), drawn from a uniform distribution (*U* [19, 24]). The initial environmental variance (*ϵ*) for species was randomly sampled from a uniform distribution *U* [0.009, 0.03], and the initial genetic variance (*V*_*i*_) was randomly sampled from a uniform distribution (*U* [0.009, 0.03]). All species started with an initial density of 1, and an extinction threshold was fixed at 10^−6^, and species falling below this density were fixed at 0. This threshold density of 10^−6^ was only used during initialisation. Additional parameter values used to simulate the communities to reach eco-evolutionary equilibrium are detailed in Table 1.

**Table 1.** Description of parameters and their values used in the study. *U* [*a, b*] is the uniform distribution between *a* and *b*.

| Symbol | Description |
| --- | --- |
| $V_i$ | Genetic variance of species $i$. |
| $h_i^2$ | Heritability of species $i$'s trait; equal to $\frac{V_i}{\rho_i}$. |
| $g_i$ | modulates the peak of temperature-dependent growth rate; fixed at 1.5 for all species. |
| $\alpha_{ii}$ | trait-independent intraspecific competition; fixed at 1 for all species. |
| $b_w$ | modulates the trade-off between temperature-dependent growth rate width, and peak; fixed at 2 for all species. |
| $\alpha_{ij}^{(A,P)}$ | pairwise inter- and intraspecific competition; intraspecific competition $\alpha_{ii}$ was fixed at 1 for all species, whereas interspecific competition $\alpha_{ij}$ was derived from equation 2 (see appendix 1). |
| $\gamma_0$ | maximum mutualistic interaction strength; fixed at 1. |
| $w$ | width of mutualistic Gaussian interaction kernel; fixed at 1. |
| $w_c$ | width of the Gaussian competition kernel, fixed at 0.2. |
| $k_i$ | mortality from the temperature tolerance curve; fixed at 0.4 for all species. |
| $T$ | Average local environmental temperature, which shifted from $22^\circ C$ to $26^\circ C$. |
| $u_i$ | Mean optimum trait values of species with units of degree Celcius. Their initial values are sampled from $U[19, 24]$. |
| $N_i^A, N_i^P$ | Population density of animal and plant species. Their initial value was 1 for all species. |
| $\epsilon$ | environmental variance, randomly sampled for all species from $U[0.009, 0.03]$. |
| $H$ | Handling time, fixed at 0.25 for all species. |

First we simulated the eco-evolutionary dynamics of plant-pollinator communities with constant genetic variance, and only the mean trait evolved in response to temperature, changes in competitive pressures and mutualistic benefits. This means that the right-hand side of Equation 7 was set to 0, keeping genetic variance constant at its initially assigned random values. This was done to let species adapt to the local environmental temperature as well as to their partners, without the erosion of genetic variance, which could happen due to stabilising selection. Thus, using Equations 5 and 6, we simulated the eco-evolutionary dynamics of plant-pollinator communities until 2 * 10^3^ time points. By this time, the plant-pollinator networks had reached a quasi-eco-evolutionary equilibrium. Subsequently, we used the final species densities and mean trait values at *t* = 2 * 10^3^ as the starting point for the press perturbation, or shift in temperature, *T*.

## 4 Shift in temperature and evolutionary response

We aimed to evaluate the distinct effects of optimum phenotype evolution and genetic variance evolution on the eco-evolutionary response of complex plant-pollinator networks, comparing these outcomes to scenarios where only mean optimum trait evolves or where neither the mean trait nor genetic variance evolves. Additionally, we sought to determine whether the strength of trait-based competition exerts a positive or negative influence on evolutionary rescue and to identify the networks most susceptible to these effects. Furthermore, we investigated how a species’ role within a network influences its eco-evolutionary response following an abrupt shift in environmental temperature, specifically examining whether being closely connected to a hyper-generalist or occupying a peripheral position within the network confers greater resilience.

We assessed the impact of an abrupt shift in temperature on the evolutionary response of species within plant-pollinator networks. Species in these networks are adapted to both their interaction partners and the local temperature. All species that had a density larger than 10^−3^ at the time of environmental shift were classified into exactly one of the following outcome classes depending on their dynamics following a shift in temperature (Fig. 1, see Fig. S11 for the exact classification rules): 1) *growth*: certain species may experience an increase in population density if these species are adapted better to the new temperature than the one before, 2) *rescue*: species initially experience a decline due to maladaptation but may eventually adapt to the new temperature and recover, exhibiting a U-shaped population trajectory. 3) *Stasis*: some species might remain stable in their population size, despite the shift in temperature 4) *decline*: certain species, due to specific eco-evolutionary factors, may decline and fail to recover, ultimately going extinct, 5) *decline-stasis*: certain species may immediately decline in population density, and then stabilise at a lower population density or just decline slowly enough so that they have not gone extinct by the end of the simulation (Fig. 1).

Thus, following the temperature shift, we quantified each of these five metrics as shown in Fig. 1 and evaluated how such metrics were impacted by three models of only mean trait evolution, both mean trait and genetic variance evolution and no evolution.

In addition, we further evaluated how species’ response to abrupt environmental change is influenced by network nestedness, network size, network connectance, modularity, species degree, species proximity to a hyper-generalist, species trait lag, and average strength of competition faced by a species before the shift in the environmental temperature. Species proximity to a hyper-generalist was measured as the minimum number of edges that leads to the hyper-generalist. The hyper-generalist is the species with the highest number of interactions. Species trait lag after the temperature shift was calculated as the difference between the new environmental temperature and species mean value of the optimum trait before the temperature shift. Connectance was quantified as the total number of possible species links divided by the product of the total number of plant and pollinator species. To assess nestedness, we employed the commonly used NODF method (Almeida-Neto *et al.*, 2008; Song *et al.*, 2017). Modularity captures how clustered a mutualistic network is. Networks with high modularity will have groups of species that have more links among group members than with species outside these groups. Network modularity was from the R package *bipartite* (Dormann *et al.*, 2009). However, these network metrics such as nestedness, connectance, network size and modularity are highly correlated (Baruah & Wittmann, 2024; Guimarães et al., 2017). We thus used Principal Component Analysis (PCA) of the four network metrics to obtain two axes of structural variation which were the first two principal components (PC1 and PC2). We then used scores of PC1 as it correlated with high nestedness and connectance and low modularity values (see Fig S1). PC2 was correlated with network size and we also evaluate the impact of network size on species eco-evolutionary response to abrupt environmental change.

To identify which explanatory variables might be significant in explaining evolutionary response based on the high-dimensional simulation data, we employed multivariate random forest regression. This method was well-suited to our analysis, as it accommodates categorical, non-linear, continuous, and transformation-invariant explanatory variables. We constructed 500 regression trees to assess the importance of these variables derived from our eco-evolutionary simulations. These regression trees are decision trees that are constructed within the random forest model. The multivariate response variable included the five outcomes we quantified: *rescue, stasis, growth*, and *decline, decline-stasis*, all of which are binomial in nature. The explanatory variables were detailed in the previous paragraph. The importance of each explanatory variable was determined by randomly shuffling out-of-bag data, that is, a part of the data that was not used to train the multivariate random forest model. Given the multivariate nature of our response variable, we utilized a Mahalanobis distance splitting rule that accounts for correlations between the different responses when constructing the regression trees. Mahalanobis distance is the statistical distance measuring how far a set of points is from a given distribution. The multivariate random forest regression was performed using the R package *RandomForestSRC* (Ishwaran & Kogalur, 2007). The multivariate random forest was used to obtain the variables which explained the most variation in the metrics that we quantified.

## 5 Results

### 5.1 Eco-evolutionary response to abrupt change in temperature: role of evolution of mean and genetic variance

We used 61 empirical plant-pollinator networks, which have connectance values ranging from 0.08 to 0.7. A sudden change in environmental temperature led to collapses of the entire plantpollinator networks, particularly when neither the mean optimum phenotype nor genetic variance can evolve, indicating that plant-pollinator networks may lack resilience to temperature shifts without any evolutionary dynamics (see example in the first row of Fig. 2). However, when the mean phenotype evolves in response to abrupt temperature change, a number of species could adapt to the new environmental conditions, leading to higher resilience in comparison to when species do not evolve (second row of Fig. 2). Furthermore, when both the mean optimum trait as well as species genetic variance was allowed to evolve we observed a much larger number of species to evolutionarily adapt to the new environmental temperature shift (third row of Fig.2).

**Figure 2.**
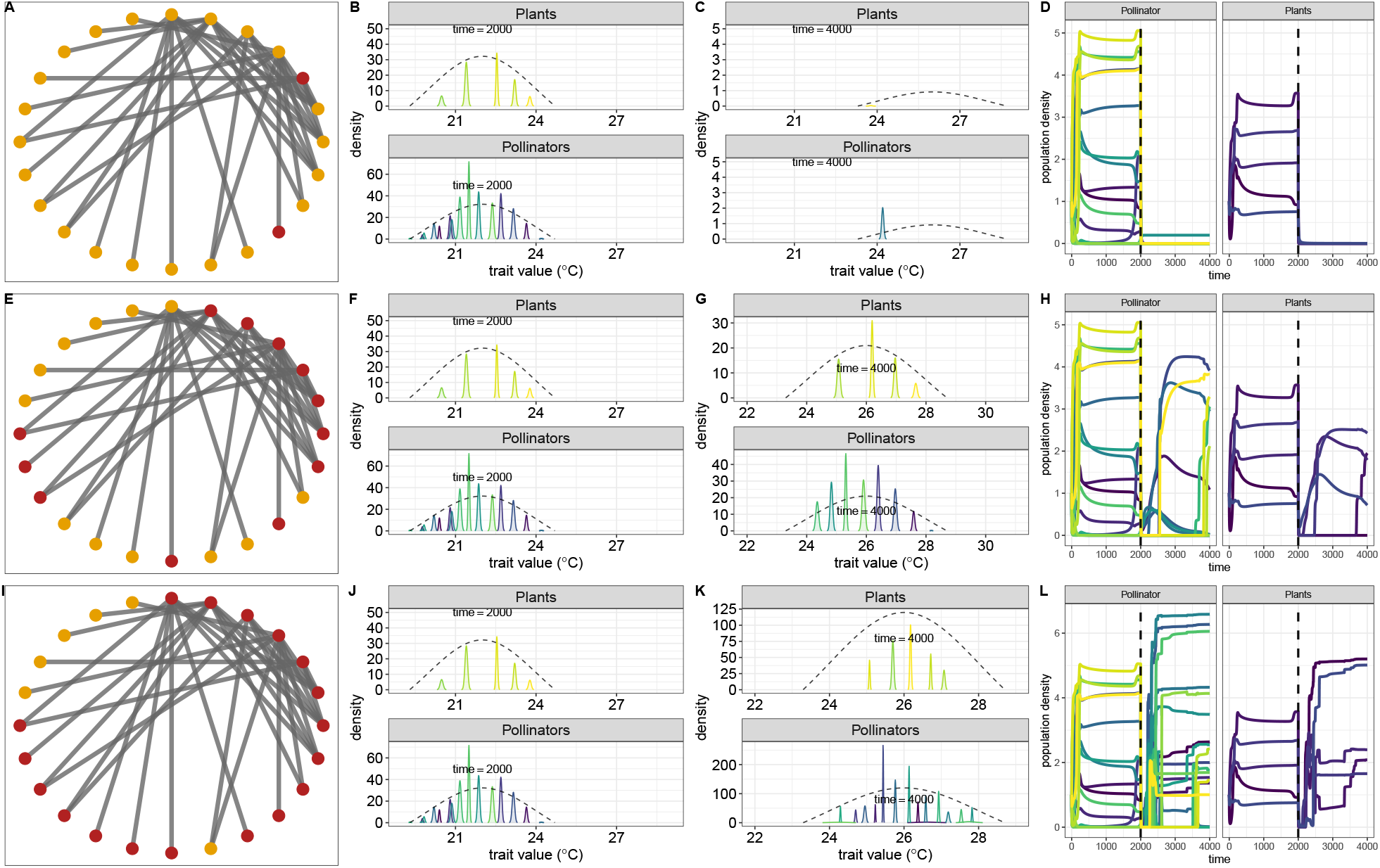
Example eco-evolutionary dynamics of a 23-species plant–pollinator network under three scenarios following a temperature shift from 22^°^C to 26^°^C at time *t* = 2000: no evolution (A–D), evolution of mean trait only (E–H), and evolution of both mean trait and genetic variance (I–L). In network panels (A, E, I), node colors indicate species that survived (dark red) or went extinct (orange). Without evolution, most species go extinct (A–D). Allowing trait evolution increases survival (E–H), and evolving both mean and variance yields the highest persistence (I–L). Dashed curves show temperature tolerance curve. Parameters follow Table 1.

When genetic variance evolves in response to a shift in the temperature, both the mean trait value of species and the constraints imposed by a fixed genetic variance are altered. This allows genetic variance to either increase or decrease in response to environmental changes. In this scenario, a large temperature shift induces a selection pressure to adapt to the new temperature, while also maintaining trait matching with mutualistic partners. As a result, for instance as shown in Fig 3, just after the environmental temperature shift there are multiple regions on the trait axis where there are two peaks of high growth rates separated by a valley of very low growth rates, indicating the transient emergence of disruptive selection (Fig. 3). For instance in Fig. 3, just after the local temperature shifts to 26 ° C, plant and pollinator density declines considerably and the per-capita growth rate has two peaks, one with the new optimum at 26 ° C and the other dictated by the group of species around 22 °C. Due to this reason, such a fitness landscape will lead to increase or decrease in trait density in the two regions of the trait axis. Due to our model formulation and random mating, trait distributions will remain normal, but trait variance can increase (see example figure S10). Eventually, this increase in trait variance will allow for faster evolution of the mean trait of these species. Consequently, adaptation to the new environmental temperature occurs faster. Once the species adapt to the new environmental temperature, stabilising selection on the mean phenotype causes species to lose the extra genetic variance they gained (Fig. 3 E-F), and they are only left with environmental variance given by *ϵ*.

**Figure 3.**
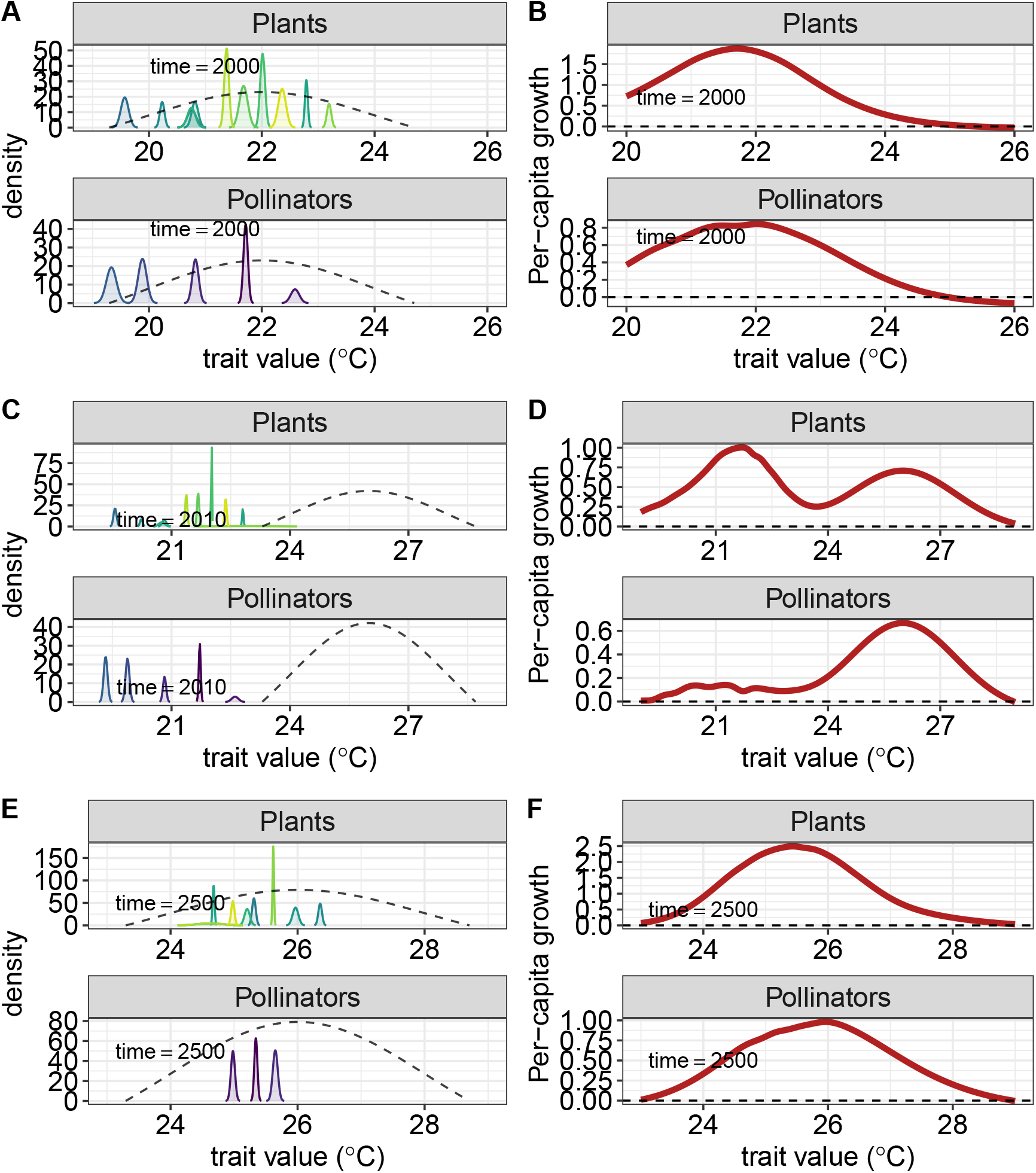
Evolution of trait distributions following an abrupt temperature shift. (A) Before the shift, species traits and growth rates peak around 22^°^C, with high-density species having more interactions; the dashed curve shows the temperature tolerance. (B) The corresponding fitness landscape, or per-capita growth rate for both plant and animal species trait values peaks around 22 ° C (this fitness curve is obtained from equation 1). (C–D) After shifting to 26^°^C, densities drop and the fitness landscape becomes bimodal at *t* = 2010 (D), reflecting opposing selection pressures: one peak at 26^°^C and another near 22^°^*C* due to the hyper-generalists, which transiently increases phenotypic variance. (E–F) By time *t* = 2500, species adapt to the new temperature, and stabilizing selection narrows the fitness peak at 26^°^ C, reducing genetic variance to baseline environmental variance.

### 5.2 Impact of competition, trait-lag, and species interaction

From the analysis of our multivariate random forest model, we found that species competition, trait-lag, species degree, and minimum path to super-generalists are important metrics explaining our multivariate binomial response (Fig. S6). We thus present our results of all the metrics below in detail.

Without evolutionary dynamics of either the mean optimum trait or the genetic variance, the number of species surviving a large shift in environmental temperature in the presence of moderate competition width *ω*_*c*_ *>* 0.1 is less than 25% overall (Fig. S12 D-H). Since, *ω*_*c*_ controls the width of species competition, higher values would indicate higher strength of competition. With evolutionary dynamics in mean trait only and large temperature shift *>* 7, the proportion of species that survived is less than when both the mean and genetic variance evolve. This was more evident in one plant-pollinator network (Fig. S12 E) than in the other (Fig. S12 A). Overall at low competition strength *ω*_*c*_ *<* 0.2, even at large temperature shift *>* 8 more than 60% of species survive when both the mean and the genetic variance evolves in comparison to only the mean trait evolving in response to large temperature shift. A larger shift in temperature may carve out a well-defined fitness peak, allowing directional selection to efficiently guide species towards the new environmental temperature. In contrast, more small shifts in temperature can lead to a smaller fitness peak in the fitness landscape which could be in addition to other shallow suboptimal local fitness peaks, where populations can become trapped and unable to climb towards higher fitness.

We found that when there is no evolution, the proportion of species that survive (have densities higher than 10^−4^ at the end of a simulation) and the proportion of species exhibiting rescue were the lowest when compared with cases where the mean optimum trait evolves, or where both the mean and the genetic variance evolve (Fig. 4A). In general, when the mean and genetic variance evolve, the proportion of species surviving is greater than when only the mean trait evolves (Fig. 4A), while the proportion of species exhibiting decline was similar when the mean optimum trait only, and mean and genetic variance of the trait evolves (Fig. 4A). In addition, the proportion of species exhibiting rescue behavior was slightly higher in the model when only the mean trait evolves in comparison to when the mean and the genetic variance evolves. Surprisingly, even when evolutionary dynamics were not at play, we also observed species exhibiting the rescue behavior (Fig. 4A, rescued). We also observed that the proportion of species exhibiting increases in species density i.e., growth, after a temperature shift of 4°*C* was lowest with evolution of just the trait mean, intermediate without evolutionary dynamics, and highest when the mean and genetic variance of the trait evolved (Fig. 4A). The proportion of species exhibiting stasis was very low (*<* 1 %) in all the three different models. Finally, the proportion of species that exhibited decline and then stasis was also high, especially when evolutionary dynamics were absent and when both mean and the genetic variance of the optimum trait evolved.

**Figure 4.**
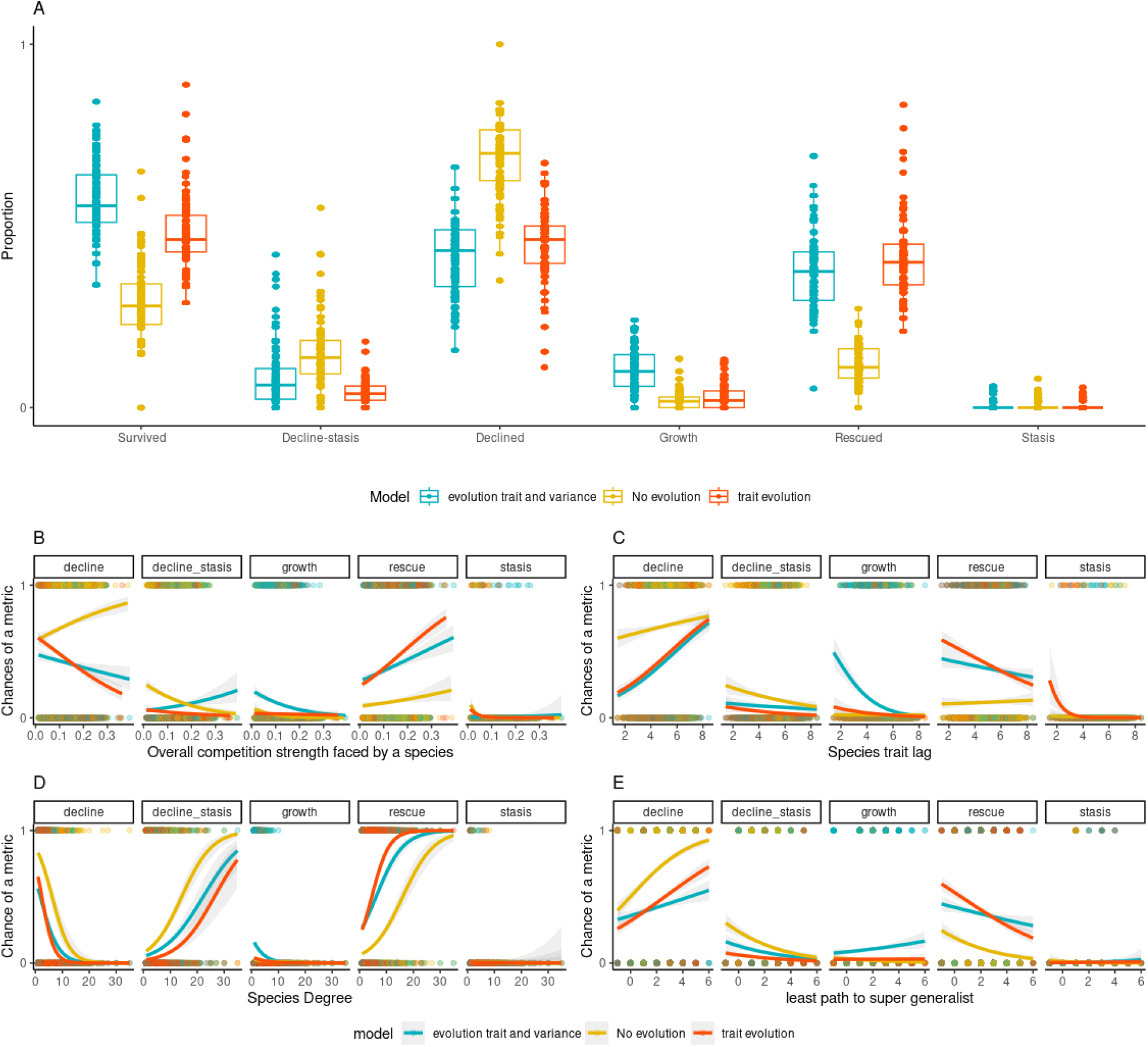
Proportion of species per network exhibiting different responses such as survival, decline, rescue, growth, stasis, or decline-stasis, after a temperature shift under three models: no evolution, evolution of mean trait only, and evolution of both mean and genetic variance. (A) Survival and growth were highest when both mean and variance evolved; decline was most frequent with no evolution. Rescue proportion were similar with mean-only and mean and genetic variance evolution. (B) Higher competition increased rescue but reduced growth, especially when variance evolved. (C) Greater trait lag led to more declines, though rescue remained possible even at high lag. (D) Species with higher degree were more likely to be rescued and less likely to decline. (E) Greater distance from a hyper-generalist increased risk of decline and reduced chances of rescue. Lines show predictions from quasi-binomial GLMs with 95% confidence intervals. Parameters as in Table 1.

In all three models, competition was a driver for species rescue (Fig. 4B). However, competition was a strong driver for species decline when there was no evolution in play. The chance of species decline at the highest competition strength was the highest when there was no evolution (Fig. 4B). Furthermore, chances of rescue were also the highest when competition was strong for all the three models. In addition, species trait lag negatively influenced the chances of rescue, species growth, and decline-stasis (Fig. 4C). Higher trait lag led to more cases of species decline in all the scenarios. As species trait lag increased, chances of evolutionary rescue declined more steeply when species mean trait evolved in comparison to when both mean and genetic variance evolved. Furthermore, only when species had mean and genetic variance evolve, there was a high chance of species density growing particularly when species had low trait lag (Fig. 4C).

We found that a species’ position within the network influences its chances of undergoing evolutionary rescue or facing decline in response to an abrupt temperature shift. Specifically, species with a higher degree are less prone to decline and have a greater chance of exhibiting evolutionary rescue, regardless of whether only the mean trait evolves or both the mean and genetic variance evolve, or when there is no evolution at all (Fig. 4D). Even when there was no evolution, we still observed species exhibiting rescue, particularly those with a high degree as well as those with a low trait-lag (Fig. 4C-D and Fig. S8). This is because, having a higher species degree maintains positive fitness despite a shift in the environmental temperature. Similarly, a low trait lag also ensures that species suffer less after a shift in the environmental temperature, which can be visually observed in Fig. S7-S8 and Fig. 4, and thus the potential for further growth increases. Furthermore, we also observed that species with a higher number of interactions also more often exhibit decline and then stasis after a sudden change in the environment. Finally, a species proximity to a super-generalist species can also reduce chances of decline as well as increase rescue chances (Fig. 4 E).

### 5.3 Impact of network structure on metrics of eco-evolutionary response

Since network architecture metrics like nestedness, modularity, connectance, and network size are correlated, we used PCA to reduce them to two axes of variation. PC1 and PC2 together explained 99% of the variation, with PC1 being positively correlated with nestedness and connectance and negatively with modularity, while network size was negatively correlated with PC2 (Fig. S1). We found that the proportion of species undergoing evolutionary rescue was positively associated with PC1 (Fig. 5B), more so when species evolved both in mean as well as in trait variance in comparison to when there was no evolution. The proportion of species that exhibited growth following an abrupt environmental temperature shift was higher when species evolved in both trait mean and genetic variance but showed slight negative association with PC1. However, the proportion of species that declined was negatively correlated with PC1, indicating that higher nestedness and connectance could lead to fewer species declines in response to abrupt temperature shifts (Fig. 5A). Finally, the fraction of species that declined and then remained stable (decline-stasis) correlated positively with PC1 when species evolved in the mean trait only (Fig. 5D). This was similarly observed when there was no evolutionary dynamics were at play. When the mean trait and genetic variance evolved the fraction of species declining and then remaining stable was very low and slightly positively associate with PC1 (Fig. 5D).

**Figure 5.**
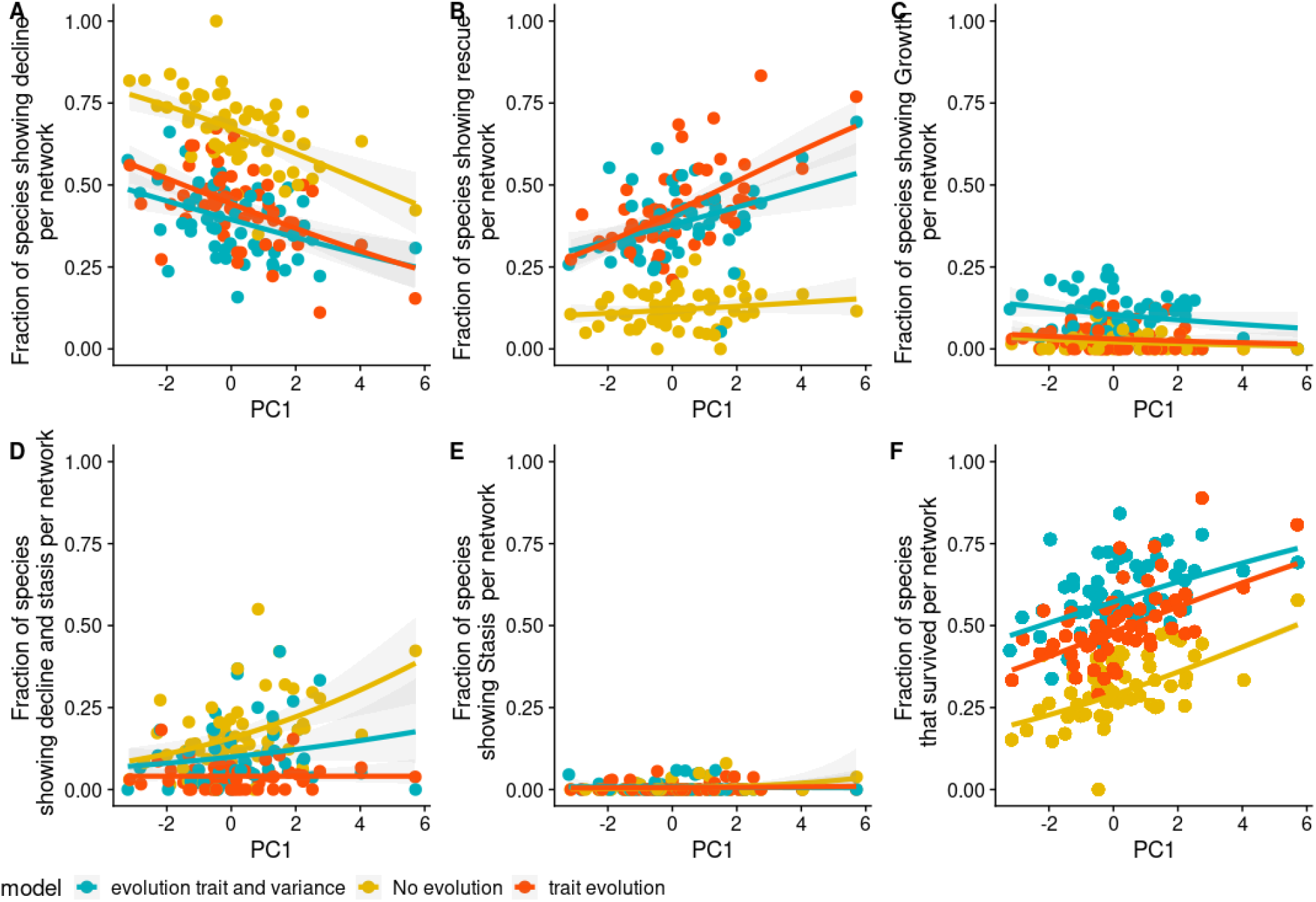
Relationship between proportion of species showing (A) decline, (B) rescue, (C) growth, (D) decline-stasis, (E) stasis only, and (F) survival in relation to PC1 which correlates with high nestedness and connectance and low modularity of plant-pollinator networks. Overall, larger PC1 values lead to more species surviving and exhibiting rescue and slightly fewer species exhibiting decline for models which had evolution of mean trait and evolution of mean trait and genetic variance. Shaded regions around lines represent 95% confidence interval of a generalised linear model with quasi-binomial error distribution. Parameters as in table 1.

## 6 Discussion

When environmental temperatures shift suddenly, species can respond in several ways: they may decline in numbers, increase in numbers, remain stable, be “rescued” through evolutionary adaptation, or decline and then remain stable (Fig.1). Whether a species can survive such temperature shifts depends on whether it can maintain positive population density through growth, stability, or adaptation via evolutionary rescue. A critical factor in species adaptation to changes in temperature is genetic variance, as higher genetic variance enables quicker adaptation to novel environmental conditions (Baruah *et al.*, 2019; Lande, 2009; Reed *et al.*, 2010). In response to environmental change, species could often respond in different ways, including dynamical changes in mean quantitative traits and genetic variance, as new selection pressures could favor phenotypes suited to the new environment. Our findings show that when genetic variance was allowed to evolve, more species successfully adapted to novel conditions and survived in comparison to scenarios where genetic variance remained fixed (Fig. 2-4).

After a sudden change in the environmental temperature, selection on the species’ traits is generally directional because their optimum traits are far way from the new environmental temperature. Such directional selection is thought to reduce genetic variance (Lynch & Walsh, 1998; Roughgarden, 1972; Taper & Chase, 1985), and thus should potentially hinder further adaptation of species in such networks. However, in most species we observed an increase in genetic variance, also known as “evolvability” (Houle, 1992; Opedal, 2019). This increase in genetic variance was largely due to a combination of different factors including complex interspecies interactions. For example, before the environmental temperature shifted to a new value, stabilizing selection maintained a single optimal trait peak, which could reduce the genetic variance of species in a plant-pollinator network (Fig. 3). In our study, however, we ensured that genetic variance did not evolve prior to the environmental shift. This was done to enable species to adapt to their partners while also aligning with the local environmental temperature, as well to ensure that stabilising selection does not erode genetic variance. Following the environmental temperature shift, some species exhibited substantial increases in genetic variance. Species that exhibited increased genetic variance tended to have higher degree, were somewhat likely to be linked to a hyper-generalist species, and were also positively associated with larger trait lags and more intense interspecific competition (Fig. S4).

Immediately after the environmental shift, there is a transient phase in the evolutionary dynamics, characterized by a lag in the response of species, as shown in Fig. 3. During this phase, certain groups of plants or pollinators initially resist shifting their mean trait values. This resistance arises because their positive growth rates are sustained by their partners, who also remain near the same trait region. At the same time, the new environmental temperature creates an additional fitness peak, exerting strong selective pressure for species to shift toward it. The opposing selection pressures—one to remain at their current trait values due to their partner species, and another to adapt to the new environmental peak—result in transient disruptive selection, leading to an increase in genetic variance. The higher the trait lag, the higher the selection pressure, and thus the higher the increase in genetic variance (Fig. S4). This, however, happens very transiently. This transient increase in genetic variance does not guarantee survival as species could already have low equilibrium density (due to species being specialist for instance) before the environment shifted, or due to strong competition which could drive them to extinction despite strong selection pressure to increase genetic variance (Fig. 3D). Eventually, the shift toward the new environmental temperature became stronger, resulting in a return to a single peak in the per-capita growth rate as more species reached the new equilibrium (Fig. 3F).

We observed some unexpected patterns in species’ rescue, growth, and decline-stasis dynamics following a shift in the environment. Notably, in the absence of evolutionary dynamics, a surprisingly high proportion of species exhibited these responses (Fig. 4A). Without evolution in either mean trait or genetic variance, many high-density species experienced an immediate decline followed by stabilization as the environment shifted. These species tended to have the lowest trait lag and the highest number of interactions (Fig. S8, which presents a clearer version of these results focused on the no-evolution scenario). Interestingly, even without evolution, some species showed growth—specifically (Fig. 4A, S7, S8). As shown in Fig. S7 as an example, these growing species low trait-lag, and after the temperature shift, they aligned more closely with the new optimum, resulting in reduced trait lag and increases in density.

The lag in species traits influenced whether species could survive the shift to a new environmental temperature, as shown in Figure 4C. When species experienced a greater trait lag, they could not adapt and match the new temperature. With evolutionary dynamics occurring in mean trait and genetic variance, species retained a larger chance of evolutionary rescue and thus survival even under maximum trait lag, in comparison with scenarios where only the mean phenotype evolved (Fig. 4C). Thus adaptation in mean and genetic variance led to a higher overall survival among species across all plant-pollinator networks (Fig. 4A). The likelihood of evolutionary rescue was also greater for species with a larger number of interactions, such as hyper-generalists, or for species that interact with such generalists (Fig. 4E). In addition, interaction with hyper-generalist also led to a crucial role in mutualistic systems not only due to the role they play in increasing community robustness against external perturbation but also their critical role in aiding recovery when communities transition to less favorable states, as observed in both empirical and theoretical studies (Baruah, 2022; Baruah & Wittmann, 2024; Bascompte & Jordano, 2013; Bascompte & Scheffer, 2023; Kaiser-Bunbury *et al.*, 2017, 2014; Lever *et al.*, 2020).

However, specialist species can also survive abrupt temperature changes, contingent on whether their genetic variance is subject to evolutionary change. In Fig. S4, we observe a substantial amount of variation in the response of specialist species and their maximum increases in genetic variance (Fig. S4B for degree < 4). This suggests that there are specialist species which still could potentially survive by means of evolution in their mean and their genetic variance. The variation in maximum increases in genetic variance for species having low number of interactions aka specialist species (i.e., degree <4) can be explained by their trait-lag rather than by the competition they faced (see Fig. S9A-C). Furthermore, overall our results reveal that when species evolve in mean trait and genetic variance, increases in genetic variance can be explained by a host of factors including the degree of species, trait lag, and competition species faced, all of these factors leading to an overall increase in genetic variance. As a consequence, as species’ genetic variance rises, species exhibit enhanced evolutionary adaptability that manifests in overall survival of species.

However, we also observed that maximum genetic variance exhibited by a species was also positively associated with chances of population decline-stasis (Fig. S4A). This pattern arose because increases in genetic variance was also suggested to be linked to species competition, (Barabás *et al.*, 2022) and trait lag here (Fig. S4D-E). However, trait-lag and species competition was also responsible for species decline, and decline-stasis in general (Fig. 4B-C). On one hand, higher increase in genetic variance facilitates rapid evolutionary changes and overall survival, but on the other hand, the conditions driving such an increase in genetic variance— significant trait lag — simultaneously heighten the chances of decline of species. Nevertheless, we believe evolution of genetic variance is on average beneficial for species survival as without it species have a higher likelihood to decline as we observe in figure 4A.

Previous studies have demonstrated that complex network architecture significantly influences the feasibility (Arroyo-Correa *et al.*, 2023; Baruah, 2022; Bastolla et al., 2009), resilience, and stability of mutualistic networks (Baruah & Wittmann, 2024; Saavedra et al., 2017). Moreover, evolutionary dynamics interact with network architecture to determine when complex mutualistic networks collapse under increasing temperatures (Baruah & Lakämper, 2024). Our study contributes to this growing body of literature by exploring how network architecture affects various aspects of species’ evolutionary responses to abrupt temperature changes. We found that network features such as nestedness and connectance positively influence the likelihood of evolutionary rescue, ultimately increasing the proportion of species that survive. Thus, in the event of an abrupt increase in temperature, if a hyper-generalist was able to adapt to a novel environmental condition, it is likely going to translate into survival of almost all other species that are interacting with such a hyper-generalist species. However, in a less nested network, there are many different species which could have similar number of interactions, and survival of many different species will be interlinked with survival of other species. Similarly, networks with high connectance which quantifies inter-guild i.e. plant-animal interactions—showed a positive correlation between evolutionary rescue and species survival (Fig. 5B-5F). This is because a higher number of inter-guild interactions i.e., higher connectance, means more positive mutualistic interactions that consequently fosters a higher number of species surviving after a temperature shift (Baruah, 2022).

Conversely, modularity, which reflects the presence of distinct species clusters within a network, was negatively correlated with evolutionary rescue (Fig. 5B). This aligns with prior findings showing that modularity can be beneficial under certain environmental conditions (Bascompte & Stouffer, 2009; Garay-Narváez et al., 2014; Olesen et al., 2007) but detrimental under others (Gilarranz *et al.*, 2017). For example, in highly modular networks, localized perturbations—such as the extinction of a species—are less likely to propagate beyond the affected module (Gilarranz *et al.*, 2017). In our study, modularity was detrimental to evolutionary rescue and overall survival. This is likely because the temperature shift impacted all species in the network uniformly, rather than being localized. Under such conditions, evolutionary shifts in mean trait of species may be constrained, as localized clusters of mutualistic interactions can hinder species from rapidly adapting to new environmental temperatures.

A previous theoretical study proposed that mutualistic interactions could reduce the likelihood of evolutionary rescue in two-species mutualistic systems (Goldberg & Friedman, 2021). This is attributed to their mutual dependence, which, combined with competition, could increase the risk of extinction rather than facilitating evolutionary rescue. Mutualistic systems, such as plant-pollinator networks and those we modeled, ranged in complexity from as few as eight species to as many as seventy. Network size, however, is not an indicative parameter to accurately capture the strength of mutualistic systems as it captures within-guild competition in addition to between-guild positive interactions. Connectance, which quantifies the number of inter-guild mutualistic interactions (or nestedness), is a true indicator of mutualistic systems in a way that as connectance between inter-guild plants and pollinators increases, so does the overall strength of positive interactions in such systems. Indeed, as connectance and nestedness increase, the proportion of species exhibiting evolutionary rescue, and survival increases (Fig. 5C, 5F). Consequently, our study does not support the notion that mutualistic interactions hinder evolutionary rescue (Goldberg & Friedman, 2021). On the contrary, they appear to promote it, particularly when species evolve along the mean and the genetic variance, and as the number of species in such a system increases. Genetic variance increased due to interspecific competition and trait lag, thereby enhancing the likelihood of evolutionary rescue. The key difference explaining why our results are different compared to those of Goldberg & Friedman (2021) is probably that the Goldberg & Friedman (2021) system comprised of only two species which were dependent on each other both in terms of mutualistic interaction and also competition. Additionally, in our model, evolution occurred through standing genetic variation, whereas in their model, populations had to wait for beneficial mutations to arise—making it unlikely that such mutations would occur in both co-dependent species within a short time frame.

Our modeling framework has several limitations. We did not include mutation rates(Wahl & Campos, 2023), which could increase genetic variance (evolvability) and aid adaptation via cryptic variation (Bell & Gonzalez, 2009). We also excluded trait plasticity, a key early response that may accelerate later adaptation (Reed *et al.*, 2010). Demographic stochasticity and environmental fluctuations were not modeled, though both can affect genetic variation, extinction risk, and adaptation, especially in small populations (Draghi *et al.*, 2024; Peniston et al., 2020). The effects of such demographic or environmental stochasticity may either support or hinder evolutionary responses. We did not model these in our framework and thus they represent avenues for future research.

## 7 Data Availability

The data will be uploaded to Zenodo. R scripts for simulations are available in the repository https://github.com/GauravKBaruah/evorescue.

## References

Alekseeva, E., Doebeli, M. & Ispolatov, I. (2020) Evolutionary adaptation of high-diversity communities to changing environments. Ecology and Evolution 10, 11941–11953, _eprint: https://onlinelibrary.wiley.com/doi/pdf/10.1002/ece3.6695.

Almeida-Neto, M., Guimarães, P., Guimarães, P.R., Loyola, R.D. & Ulrich, W. (2008) A consistent metric for nestedness analysis in ecological systems: reconciling concept and measurement. Oikos 117, 1227–1239.

Amarasekare, P. & Johnson, C. (2017) Evolution of Thermal Reaction Norms in Seasonally Varying Environments. The American Naturalist 189, E31–E45.

Arroyo-Correa, B., Jordano, P. & Bartomeus, I. (2023) Intraspecific variation in species interactions promotes the feasibility of mutualistic assemblages. Ecology Letters 26, 448–459,_eprint: https://onlinelibrary.wiley.com/doi/pdf/10.1111/ele.14163.

Barabás, G., Parent, C., Kraemer, A., Van de Perre, F. & De Laender, F. (2022) The evolution of trait variance creates a tension between species diversity and functional diversity. Nature Communications 13, 2521, number: 1.

Baruah, G. (2022) The impact of individual variation on abrupt collapses in mutualistic networks. Ecology Letters 25, 26–37, _eprint: https://onlinelibrary.wiley.com/doi/pdf/10.1111/ele.13895.

Baruah, G., Clements, C.F., Guillaume, F. & Ozgul, A. (2019) When Do Shifts in Trait Dynamics Precede Population Declines? The American Naturalist 193, 633–644.

Baruah, G., Clements, C.F. & Ozgul, A. (2021) Effect of habitat quality and phenotypic variation on abundance- and trait-based early warning signals of population collapses. Oikos 130, 850–862.

Baruah, G. & Lakämper, T. (2024) Stability, resilience and eco-evolutionary feedbacks of mutualistic networks to rising temperature. Journal of Animal Ecology n/a, _eprint: https://onlinelibrary.wiley.com/doi/pdf/10.1111/1365-2656.14118.

Baruah, G. & Wittmann, M. (2024) Reviving collapsed plant–pollinator networks from a single species. PLOS Biology 22, e3002826.

Bascompte, J. & Jordano, P. (2013) Mutualistic networks. Princeton University Press.

Bascompte, J., Jordano, P., Melián, C.J. & Olesen, J.M. (2003) The nested assembly of plant–animal mutualistic networks. Proceedings of the National Academy of Sciences 100, 9383–9387.

Bascompte, J. & Scheffer, M. (2023) The Resilience of Plant–Pollinator Networks. Annual Review of Entomology 68, 363–380, _eprint: 10.1146/annurev-ento-120120-102424.

Bascompte, J. & Stouffer, D.B. (2009) The assembly and disassembly of ecological networks. Philosophical Transactions of the Royal Society B: Biological Sciences 364, 1781–1787.

Bastolla, U., Fortuna, M.A., Pascual-García, A., Ferrera, A., Luque, B. & Bascompte, J. (2009) The architecture of mutualistic networks minimizes competition and increases biodiversity. Nature 458, 1018–1020, number: 7241.

Bell, G. & Gonzalez, A. (2009) Evolutionary rescue can prevent extinction following environmental change. Ecology Letters 12, 942–948.

Bolchoun, L., Drossel, B. & Allhoff, K.T. (2017) Spatial topologies affect local food web structure and diversity in evolutionary metacommunities. Scientific Reports 7, 1818.

Cantwell-Jones, A., Tylianakis, J.M., Larson, K. & Gill, R.J. (2024) Using individual-based trait frequency distributions to forecast plant-pollinator network responses to environmental change. Ecology Letters 27, e14368, _eprint: https://onlinelibrary.wiley.com/doi/pdf/10.1111/ele.14368.

Chevin, L.M., Lande, R. & Mace, G.M. (2010) Adaptation, plasticity and extintion in a changing enviroment: towards a predictive theory. PLoS biology 8, 1–8.

Dormann, C.F., Fruend, J., Bluethgen, N. & Gruber, B. (2009) Indices, graphs and null models: analyzing bipartite ecological networks. The Open Ecology Journal 2, 7–24.

Draghi, J.A., McGlothlin, J.W. & Kindsvater, H.K. (2024) Demographic feedbacks during evolutionary rescue can slow or speed adaptive evolution. Proceedings of the Royal Society B: Biological Sciences 291, 20231553.

Duchenne, F., Thébault, E., Michez, D., Elias, M., Drake, M., Persson, M., Rousseau-Piot, J.S., Pollet, M., Vanormelingen, P. & Fontaine, C. (2020) Phenological shifts alter the seasonal structure of pollinator assemblages in Europe. Nature Ecology & Evolution 4, 115–121.

Freimuth, J., Bossdorf, O., Scheepens, J.F. & Willems, F.M. (2022) Climate warming changes synchrony of plants and pollinators. Proceedings of the Royal Society B: Biological Sciences 289, 20212142.

Garay-Narváez, L., Flores, J.D., Arim, M. & Ramos-Jiliberto, R. (2014) Food web modularity and biodiversity promote species persistence in polluted environments. Oikos 123, 583–588,_eprint: https://onlinelibrary.wiley.com/doi/pdf/10.1111/j.1600-0706.2013.00764.x.

Gilarranz, L.J., Rayfield, B., Liñán-Cembrano, G., Bascompte, J. & Gonzalez, A. (2017) Effects of network modularity on the spread of perturbation impact in experimental metapopulations. Science 357, 199–201.

Goldberg, Y. & Friedman, J. (2021) Positive interactions within and between populations decrease the likelihood of evolutionary rescue. PLOS Computational Biology 17, e1008732.

Guimarães, P.R., Jordano, P. & Thompson, J.N. (2011) Evolution and coevolution in mutualistic networks. Ecology Letters 14, 877–885.

Guimarães, P.R., Pires, M.M., Jordano, P., Bascompte, J. & Thompson, J.N. (2017) Indirect effects drive coevolution in mutualistic networks. Nature 550, 511–514.

Hegland, S.J., Nielsen, A., Lázaro, A., Bjerknes, A.L. & Totland, Ø. (2009) How does climate warming affect plant-pollinator interactions? Ecology Letters 12, 184–195, _eprint: https://onlinelibrary.wiley.com/doi/pdf/10.1111/j.1461-0248.2008.01269.x.

Houle, D. (1992) Comparing evolvability and variability of quantitative traits. Genetics 130, 195–204.

Ishwaran, H. & Kogalur, U. (2007) Random survival forests for r. R News 7, 25–31.

Kaiser-Bunbury, C.N., Mougal, J., Whittington, A.E., Valentin, T., Gabriel, R., Olesen, J.M. & Blüthgen, N. (2017) Ecosystem restoration strengthens pollination network resilience and function. Nature 542, 223–227, number: 7640.

Kaiser-Bunbury, C.N., Vázquez, D.P., Stang, M. & Ghazoul, J. (2014) Determinants of the microstructure of plant–pollinator networks. Ecology 95, 3314–3324.

Klausmeier, C.A., Osmond, M.M., Kremer, C.T. & Litchman, E. (2020) Ecological limits to evolutionary rescue. Philosophical Transactions of the Royal Society B: Biological Sciences 375, 20190453.

Labonté, A., Monticelli, L.S., Turpin, M., Felten, E., Laurent, E., Matejicek, A., Biju-Duval, L., Ducourtieux, C., Vieren, E., Deytieux, V., Cordeau, S., Bohan, D. & Vanbergen, A.J. (2023) Individual flowering phenology shapes plant–pollinator interactions across ecological scales affecting plant reproduction. Ecology and Evolution 13, e9707, _eprint: https://onlinelibrary.wiley.com/doi/pdf/10.1002/ece3.9707.

Lande, R. (2009) Adaptation to an extraordinary environment by evolution of phenotypic plasticity and genetic assimilation. Journal of Evolutionary Biology 22, 1435–1446.

Lever, J.J., van de Leemput, I.A., Weinans, E., Quax, R., Dakos, V., van Nes, E.H., Bascompte, J. & Scheffer, M. (2020) Foreseeing the future of mutualistic communities beyond collapse. Ecology Letters 23, 2–15.

Lever, J.J., van Nes, E.H., Scheffer, M. & Bascompte, J. (2014) The sudden collapse of pollinator communities. Ecology Letters.

Lynch, M. & Walsh, B. (1998) Genetics and analysis of quantitative traits.

Menzel, A., Sparks, T.H., Estrella, N., Koch, E., Aasa, A., Ahas, R., Alm-Kübler, K., Bissolli, P., Braslavská, O., Briede, A., Chmielewski, F.M., Crepinsek, Z., Curnel, Y., Dahl, Å., Defila, C., Donnelly, A., Filella, Y., Jatczak, K., Måge, F., Mestre, A., Nordli, Ø., Peñuelas, J., Pirinen, P., Remišová, V., Scheifinger, H., Striz, M., Susnik, A., Van Vliet, A.J.H., Wielgolaski, F.E., Zach, S. & Zust, A. (2006) European phenological response to climate change matches the warming pattern. Global Change Biology 12, 1969–1976, _eprint: https://onlinelibrary.wiley.com/doi/pdf/10.1111/j.1365-2486.2006.01193.x.

Miller-Rushing, A.J. & Primack, R.B. (2008) Global Warming and Flowering Times in Thoreau’s Concord: A Community Perspective. Ecology 89, 332–341, _eprint: https://onlinelibrary.wiley.com/doi/pdf/10.1890/07-0068.1.

Morgan, R., Finnøen, M.H., Jensen, H., Pélabon, C. & Jutfelt, F. (2020) Low potential for evolutionary rescue from climate change in a tropical fish. Proceedings of the National Academy of Sciences 117, 33365–33372.

Olesen, J.M., Bascompte, J., Dupont, Y.L. & Jordano, P. (2007) The modularity of pollination networks. Proceedings of the National Academy of Sciences 104, 19891–19896.

Opedal, Ø.H. (2019) The evolvability of animal-pollinated flowers: towards predicting adaptation to novel pollinator communities. New Phytologist 221, 1128–1135, _eprint: https://onlinelibrary.wiley.com/doi/pdf/10.1111/nph.15403.

Peniston, J.H., Barfield, M., Gonzalez, A. & Holt, R.D. (2020) Environmental fluctuations can promote evolutionary rescue in high-extinction-risk scenarios. Proceedings of the Royal Society B: Biological Sciences 287, 20201144.

Reed, T.E., Waples, R.S., Schindler, D.E., Hard, J.J. & Kinnison, M.T. (2010) Phenotypic plasticity and population viability: the importance of environmental predictability. Proceedings. Biological sciences / The Royal Society 277, 3391–400.

Roughgarden, J. (1972) Evolution of Niche Width. The American Naturalist 106, 683–718.

Saavedra, S., Rohr, R.P., Bascompte, J., Godoy, O., Kraft, N.J.B. & Levine, J.M. (2017) A structural approach for understanding multispecies coexistence. Ecological Monographs 87, 470–486.

Scranton, K. & Amarasekare, P. (2017) Predicting phenological shifts in a changing climate. Proceedings of the National Academy of Sciences 114, 13212–13217.

Song, C., Rohr, R.P. & Saavedra, S. (2017) Why are some plant–pollinator networks more nested than others? Journal of Animal Ecology 86, 1417–1424.

Taper, M.L. & Chase, T.J. (1985) Quantitative Genetic Models for the Coevolution of Character Displacement. Ecology 66, 355–371, _eprint: https://onlinelibrary.wiley.com/doi/pdf/10.2307/1940385.

Thompson, J.N., Nuismer, S.L. & Gomulkiewicz, R. (2002) Coevolution and Maladaptation1. Integrative and Comparative Biology 42, 381–387.

Trunschke, J., Junker, R.R., Kudo, G., Alexander, J.M., Richman, S.K. & Till-Bottraud, I. (2024) Effects of climate change on plant-pollinator interactions and its multitrophic consequences. Alpine Botany.

Wahl, L.M. & Campos, P.R.A. (2023) Evolutionary rescue on genotypic fitness landscapes. Journal of The Royal Society Interface 20, 20230424.

Yamamichi, M. & Miner, B.E. (2015) Indirect evolutionary rescue: prey adapts, predator avoids extinction. Evolutionary Applications 8, 787–795, _eprint: https://onlinelibrary.wiley.com/doi/pdf/10.1111/eva.12295.

Åkesson, A., Curtsdotter, A., Eklöf, A., Ebenman, B., Norberg, J. & Barabás, G. (2021) The importance of species interactions in eco-evolutionary community dynamics under climate change. Nature Communications 12, 4759, number: 1.

