## appendix for "Evolution of genetic variance and its consequences for eco-evolutionary responses in complex mutualistic networks"

### 1 Mutualistic eco-evolutionary model

#### 1.1 Quantitative Genetics and Lotka-Volterra dynamics

We model the dynamics of pollinators and plants in an ecologically relevant quantitative trait $z$. Each individual belonging to a guild of pollinators or plants can be described with its trait $z$, which varies among individuals within a species. Now the number of individuals within species $i$ at time $t$ for pollinators (animals) will be $N_i^A(t)$ and for the plants it will be $N_i^P(t)$, and the distribution of their traits within each species $i$ at time $t$ can be given by a function $p_i^{(A,P)}(z,t)$ such that

$$\int p_i^{(A,P)}(z,t)dz = 1 \tag{S1}$$

at every time $t$; where the integral over the trait axis goes from minus infinity to plus infinity.

We work in the quantitative genetic limit, i.e., the trait in question is determined by many independent loci with small additive effects and under the weak selection assumption. Due to this, the trait is normally distributed (Falconer & Mackay 1996). Thus, the trait distribution can be written as:

$$p_i^{(A,P)}(z,t) = \frac{1}{\sqrt{2\pi(\rho_i^{(A,P)})^2}} \exp \frac{-(z - u_i^{(A,P)}(t))^2}{2(\rho_i^{(A,P)})^2},$$

where $u_i^{(A,P)}$ is the mean trait value for the species $i$ (could be plants or pollinators) and $(\rho_i^{(A,P)})^2$ is the trait variance or phenotypic variance. Here, we assume there is no plasticity in the trait, but there is environmental variance that contributes to the phenotypic variance. Thus, the trait variance $(\rho_i^{(A,P)})^2$ is the combination of genetic variance and environmental variance. The governing dynamical equations of population dynamics can be written with slightly modified Lotka-Volterra equations. Following Barabas & D'Andrea (2016) and Baruah (2022), the per-capita growth rate of pollinator species $i$ can then be written as

$$\begin{aligned} r_i^A(z) = \underbrace{b(z,T)}_{\text{growth rate}} - \underbrace{\alpha_{ii}^A N_i^A}_{\text{intraspecific competition}} - \underbrace{\int \sum_{j=1, i\neq j}^{S_A} N_j^A \alpha(z,z''') p_j^A(z''',t) dz'''}_{\text{trait-based interspecific competition}} + \\ \underbrace{\int \frac{\sum_{k=1}^{S_P} \gamma(z,z') A_{ik} N_k^P(t) p_k^P(z',t)}{1 + H \int \sum_{k=1}^{S_P} \gamma(z,z'') A_{ik} N_k^P(t) p_k^P(z'',t) dz''} dz'}_{\text{trait-based mutualistic interaction}}. \end{aligned} \tag{S2}$$

The per-capita growth rate for a plant species can be written analogously. Here $b(z,T)$ is the growth rate independent of competition or mutualistic benefits; $a_{ii}^{A,P}$ is the strength of density dependence or intraspecific competition in animals or plants; $H$ is the handling time; $\alpha(z,z''')$ is the interspecific competition kernel between individuals of phenotype $z$ and $z'''$ belonging to the same guild of species; here, $\alpha(z,z''') = \alpha_0 \exp(-\frac{(z-z''')^2}{\omega_c^2})$, $\omega_c^2$ is the width of the competition kernel; $\gamma(z,z')$ is the function that captures the mutualistic interactions among a pollinator individual with trait $z$ and a plant individual with trait $z'$. We can take this function to be a Gaussian:

$$\gamma(z,z') = \gamma_0 \exp\frac{-(z-z')^2}{w^2},$$

where $\gamma_0$ is the maximum strength of mutualistic interactions and $w^2$ is the width that controls how strongly two individuals interact. The more similar traits of two individuals belonging to two different species, namely plants and pollinators, the stronger is the mutualistic benefit. From equation S2 we can arrive at trait dynamics and population dynamics equations given in the section below.

Equation S2 represents the per-capita growth rate of an individual with phenotype $z$ interacting facilitatively with other individuals with phenotype $z'$ belonging to a species of another guild, with $p_k^{(A,P)}(z',t)$ being the distribution of the trait $z'$, and competitively with other individual phenotypes $z'''$ belonging to the same guild of species. The integration in the third and fourth term of equation S2 goes over the entire trait space and is summed for all the species belonging to a guild. Thus growth and mutualistic interactions only depend on the phenotype $z$, not on species identity. Also note that mutualistic interaction follows a type 2 functional response. If $H = 0$, mutualistic benefits follow a linear type 1 functional response.

#### 1.2 Population dynamics and trait dynamics

We can arrive at continuous population and trait dynamics (see Barabas & D'Andrea 2016, Baruah 2022, Baruah & Wittmann 2024) of species $i$ (could be represented similarly for both plants or pollinators, but shown here for pollinators) by integrating over all trait space $z$ which can be written as following (Barabas & D'Andrea 2016, Baruah 2022, Baruah & Wittmann 2024):

$$\frac{dN_i^A}{dt} = N_i^A(t)\int r_i^A(z)p_i^A(z,t)dz. \tag{S3}$$

Substituting equation S2 into S3, we get

$$\begin{aligned}\frac{dN_i^A}{dt} = N_i^A(t)\int\Bigg(b(z,T) - \alpha_{ii}^A N_i^A - \int\sum_{j=1,i\neq j}^{S_A} N_j^A\alpha(z,z''')p_j^A(z''',t)dz''' + \\ \int\frac{\sum_{k=1}^{S_P}\gamma(z,z')A_{ik}N_k^P(t)}{1+H\int\sum_{k=1}^{S_P}\gamma(z,z'')A_{ik}N_k^P(t)p_k^P(z'',t)dz''}p_k^P(z',t)dz'\Bigg)p_i^A(z,t)dz\end{aligned} \tag{S4}$$

We can further write equation S4 as:

$$\frac{dN_i^A}{dt} = N_i^A(t)\Bigg(b_i^A(T) - \alpha_{ii}^A N_i - \sum_{j\neq i}\alpha_{ij}^A N_j^A + \gamma_i\Bigg) \tag{S5}$$

with temperature-dependent growth rate

$$b_i^A(T) = \int b(z,T)p_i^A(z,t)dz = \frac{g_i}{b_w}\exp(-\frac{(T-u_i)^2}{2b_w^2+2\rho_i^2})\frac{b_w}{\sqrt{b_w^2+\rho_i^2}} - k_i, \tag{S6}$$

interspecific competition coefficient

$$\alpha_{ij}^A = \frac{\omega_c}{\sqrt{2\rho_i^2+2\rho_j^2+\omega_c^2}}\exp(\frac{-(u_i-u_j)^2}{2\rho_i^2+2\rho_j^2+\omega_c^2})$$

and cumulative mutualistic effect

$$\gamma_i = \int\int\frac{\sum_{k=1}^{S_P}\gamma(z,z')A_{ik}N_k^P(t)}{1+H\int\sum_{k=1}^{S_P}\gamma(z,z'')A_{ik}N_k^P(t)p_k^P(z'',t)dz''}p_k^P(z',t)dz'p_i^A(z,t)dz, \tag{S7}$$

which we solved numerically.

Assuming the phenotypic trait is based on the quantitative genetic limit – i.e., it is controlled by many loci each of small additive effects under weak selection – the dynamics of the mean phenotype $u_i^{(A)}(t)$ of interest, here for the pollinators, can then be written as (Baruah 2022, Barabas & D'Andrea 2016)

$$\begin{aligned}\frac{du_i^A}{dt} = h_i^2\int(z-u_i^A)\Bigg(b(z,T) - \alpha_{ii}^A N_i - \int\sum_{j=1,i\neq j}^{S_A} N_j^A\alpha(z,z''')p_j^A(z''',t)dz''' + \\ \int\frac{\sum_{k=1}^{S_P}\gamma(z,z')A_{ik}N_k^P(t)}{1+H\int\sum_{k=1}^{S_P}\gamma(z,z'')A_{ik}N_k^P(t)p_k^P(z'',t)dz''}p_k^P(z',t)dz'\Bigg)p_i^A(z,t)dz,\end{aligned} \tag{S8}$$

where $h_i^2$ is the broad sense heritability of the trait. We can write this more compactly as

$$\frac{du_i^A}{dt} = h_i^2 \left( \overline{b_i(T)} - \sum_{j=1, i \neq j}^{S_A} \overline{\alpha_{ij}}^A N_j^A + \overline{\gamma_i} \right), \tag{S9}$$

where

$$\overline{b_i(T)} = \int (z - u_i^A) b(z,T) p_i^A(z,t) dz = \frac{g_i}{b_w} \exp(-\frac{(T-u_i)^2}{2b_w^2 + 2\rho_i^2}) \frac{\rho_i^2 b_w (T - u_i)}{(b_w^2 + \rho_i^2)^{1.5}} \tag{S10}$$

represents the impact of direct selection on temperature tolerance and

$$\overline{\alpha_{ij}}^A = \frac{\rho_i \omega_c (u_j - u_i)}{(2\rho_i^2 + 2\rho_j^2 + \omega_c^2)^{1.5}} \exp(-\frac{(u_i - u_j)^2}{(2\rho_i^2 + 2\rho_j^2 + \omega_c^2)}) \tag{S11}$$

represents the impact of competition with other species in the guild on mean trait evolution. And finally

$$\overline{\gamma_i} = \int \int (z - u_i^A) \frac{\sum_{k=1}^{S_P} \gamma(z,z') A_{ik} N_k^P(t)}{1 + H \int \sum_{k=1}^{S_P} \gamma(z,z'') A_{ik} N_k^P(t) p_k^P(z'',t) dz''} p_k^P(z',t) dz' \Bigg) p_i^A(z,t) dz \tag{S12}$$

is the impact of mutualistic interactions on mean trait evolution, which is again evaluated numerically. Thus equation S4 and equation S9 represent population dynamics and the dynamics of the trait mean, respectively. In the next sections, we derive the dynamics of genetic variance and the assumptions that we make to arrive at the genetic variance dynamics.

### 2 Weak selection assumption, fitness, and phenotypic variance:

In this section, some of the derivations and assumptions are simplified and elaborated from the work of Barabás *et al.* (2022). We expand on their work and explain the derivations from the perspective of a community that involves plant-pollinator interactions.

Let us introduce some useful assumptions before introducing the governing dynamics of trait variance. The total population density of species $i$ is denoted by $N_i^{(A,P)}$ (let's assume $N_i$ as a general term for either plants or animals for now). We assume a simple life-cycle of a population with viability selection first and then reproduction after, such that $N_i(t)p_i(z,t)$ is the population density of the phenotype $z$ before selection, $\widetilde{N_i(t)}\widetilde{p_i(z,t)}$ is the population density of the phenotype $z$ after selection and finally $N_i'(t+s)p_i(z,t+s)'$ is the population density after reproduction, i.e., in the next generation, $s$ being the generation time of species $i$. In nature, viability selection and reproduction could happen separately, however, we do not distinguish between the steps, and assume it happens in the second step itself, i.e., viability selection and reproduction happen together. We felt it is good to elucidate the actual differences that could occur in nature.

If we can denote $f_i(z,t)$ as the fitness of phenotype $z$ of species $i$ then

$$f_i(z,t) p_i(z,t) N_i(t) = \widetilde{N_i(t)} \widetilde{p_i(z,t)} \tag{S13}$$

Here, $f_i(z,t)$ encodes both birth and death processes and accounts for the dependency on trait distributions of interacting species and their abundances. Thus, the role of reproduction together with viability selection would be to shift the trait distribution in terms of the mean and potentially the variance, but the distribution is assumed to remain normal. Thus, we can write

$$p_i(z,t) \neq p_i(z,t+s)'; \widetilde{N_i}(t) = N_i'(t+s). \tag{S14}$$

The dynamics of population densities from one to the next generation can be obtained by integrating equation S13 as:

$$\widetilde{N_i(t)} \int \widetilde{p_i(z,t)} dz = N_i(t) \int f_i(z,t) p_i(z,t) dz \tag{S15}$$

From equation S1, the integral on the left hand side is 1, hence, the above equation S15 can be written as:

$$\widetilde{N_i}(t) = N_i(t) \int f_i(z,t) p_i(z,t) dz. \tag{S16}$$

Since, from equation S14 we can rewrite equation S15 as

$$N_i^{'}(t+s) = N_i(t) \int f_i(z,t) p_i(z,t) dz. \tag{S17}$$

Next, the phenotypic variance after selection i.e., $\widetilde{\rho_i}(t)$ of a variable $z$, can be written as:

$$\widetilde{\rho_i}(t) = \int (z - \widetilde{u_i}(t))^2 \widetilde{p_i(z,t)} dz \tag{S18}$$

where, $\widetilde{u_i}(t)$ is the mean of the distribution $\widetilde{p_i(z,t)}$. Rearranging equation S13 we get,

$$\widetilde{p_i(z,t)} = \frac{f_i(z,t)p_i(z,t)N_i(t)}{\widetilde{N_i(t)}}. \tag{S19}$$

Substituting equation S15, we obtain:

$$\widetilde{p_i(z,t)} = \frac{f_i(z,t)p_i(z,t)N_i(t)}{N_i(t)\int f_i(z',t)p_i(z',t)dz'} = \frac{f_i(z,t)p_i(z,t)}{\int f_i(z',t)p_i(z',t)dz'}. \tag{S20}$$

Now substituting equation S20 into S18, we get

$$\widetilde{\rho_i}(t) = \frac{\int (z - \widetilde{u_i}(t))^2 f_i(z,t)p_i(z,t)dz}{\int f_i(z,t)p_i(z,t)dz}. \tag{S21}$$

Next, we apply a weak selection limit for a finite population following Bürger (2011) and Wild & Traulsen (2007), Barabas & D'Andrea (2016), Lande (1976). The key idea is to let the generation time $s$ be small so that selection per generation becomes weak. Fitness, $f_i(z,t)$, can then be written as a Taylor approximation, $f_i(z,t) = 1 + sr_i(z,t)$, where $s << 1$, and $r_i(z,t)$ is the per-capita growth rate of phenotype $z$ for species $i$. With $r_i(z,t)$ as the per-capita growth rate which as units of inverse of generation time, then selection intensity, $s$ i.e., has units of generation time, such that fitness is a unit-less quantity. $s$ is usually assumed to be small, meaning that selection is weak and changes in phenotype per generation is also very small. One of the assumptions of genetics of quantitative traits is random mating and linkage equilibrium – which means that association of alleles from different loci contributing to a quantitative is unlinked, or in other words they are inherited independently. Random mating and weak selection intensity allows for linkage equilibrium (assuming no epistasis). Thus, with weak selection assumption in such polygenic traits in quasi-equilibrium, a population evolves as if it is in linkage equilibrium see eq-39 of Nagylaki (1993). Hence, we can approximate $f_i(z,t)$ as a Taylor approximation and then equation S21 can be written as

$$\widetilde{\rho_i}(t) = \frac{\int (z - \widetilde{u_i}(t))^2 (1 + sr_i(z,t))p_i(z,t)dz}{\int (1 + sr_i(z,t))p_i(z,t)dz} \tag{S22}$$

$$= \frac{\int (z - \widetilde{u_i}(t))^2 p_i(z,t)dz + s\int (z - \widetilde{u_i}(t))^2 r_i(z,t)p_i(z,t)dz}{\int p_i(z,t)dz + s\int r_i(z,t)p_i(z,t)dz} \tag{S23}$$

Now we Taylor-expand the above function around $s = 0$ (under no selection) and then ignore higher-order terms, i.e., ,

$$\widetilde{\rho_i}(t) = \widetilde{\rho_i}(t)\Big|_{s=0} + \frac{\frac{\partial}{\partial s}\widetilde{\rho_i}(t)}{1!}\Big|_{s=0}(s - 0) + O(s^2), \tag{S24}$$

where the first term is

$$\widetilde{\rho_i}(t)(s)\Big|_{s=0} = \int (z - \widetilde{u_i}(t))^2 p_i(z,t)dz, \tag{S25}$$

and, using the quotient rule, the second term is:

$$\frac{\frac{\partial}{\partial s}\widetilde{\rho_i}(t)}{1!}\Big|_{s=0}(s - 0) = s\left[\int (z - \widetilde{u_i}(t))^2 r_i(z,t)p_i(z,t)dz - \left(\int (z - \widetilde{u_i}(t))^2 p_i(z,t)dz\right)\left(\int r_i(z,t)p_i(z,t)dz\right)\right]. \tag{S26}$$

We now need to represent $\widetilde{u_i}(t)$ in terms of $u_i(t)$. Note that $\widetilde{u_i}$ is the mean of the phenotypic trait after selection and before reproduction. Following the Barabás *et al.* (2022) formulation on mean trait after selection, we can write

$$\widetilde{u_i}(t) = u_i(t) + sq_i(u_i,t) + O(s^2) \tag{S27}$$

where, $q_i$ encodes a function i.e., $\left(\int(z-u_i(t))r_i(z)p_i(z,t)dz\right)$ of per-capita growth rate, and distribution of other interacting species and their abundances, $O(s^2)$ contains higher-order terms in $s$, $u_i$ is the mean trait before selection. Substituting equation S27 in equation S25 and equation S26 to get:

$$\widetilde{\rho_i}(t)(s)\bigg|_{s=0} = \int(z-u_i(t)-sq_i)^2 p_i(z)dz + O(s^2), \tag{S28}$$

$$= \int(z-u_i(t))^2 p_i(z,t)dz - \int 2sq_i(z-u_i(t))p_i(z,t)dz - \int s^2 q_i^2 p_i(z,t)dz + O(s^2), \tag{S29}$$

$$= \int(z-u_i(t))^2 p_i(z,t)dz + O(s^2) \tag{S30}$$

$$= \rho_i(t) + O(s^2), \tag{S31}$$

where the second term $\int 2sq_i(z-u_i(t))p_i(z,t)dz = 0$ due to the fact that $\int(z-u_i(t))p_i(z,t)dz = 0$, and similarly

$$\frac{\frac{\partial}{\partial s}\widetilde{\rho_i}(t)}{1!}\bigg|_{s=0}(s-0) = s\left[\int(z-u_i(t))^2 r_i(z,t)p_i(z,t)dz - \rho_i(t)\left(\int r_i(z,t)p_i(z,t)dz\right)\right] + O(s^2). \tag{S32}$$

Thus we can finally rewrite equation S24 after substituting the result of equation S28 and S32 as

$$\widetilde{\rho_i}(t) = \rho_i(t) + s\left[\int(z-u_i(t))^2 r_i(z,t)p_i(z,t)dz - \rho_i(t)\left(\int r_i(z,t)p_i(z,t)dz\right)\right] + O(s^2) \tag{S33}$$

$$= \rho_i(t) + s\int\left[(z-u_i(t))^2 - \rho_i(t)\right] r_i(z,t)p_i(z,t)dz + O(s^2). \tag{S34}$$

Before we finally get to the equation of genetic variance dynamics, we need to introduce some basic relationships between offspring and mid-parent phenotypes.

### 2.1 Segregation variance, mid-parent and offspring phenotype, genetic variance dynamics:

Here in this section, the aim is to get an analytical expression of total variance of offspring phenotype. Based on the law of total variance

$$Var[Y] = Var[E[Y|X]] + E[Var[Y|X]] \tag{S35}$$

the total phenotypic variance of offspring is the sum of the variance of expected offspring phenotype given the mid-parent phenotype and the residual or segregation variance, which cannot be explained by the mid-parent phenotype.

Consider $z$ as the mid-parent phenotypic value with $z = \frac{z_f+z_m}{2}$, where $z_f, z_m$ are the phenotypic values of the two parents. In that case, from classic quantitative genetics theory (Falconer & Mackay 1996), the expected offspring phenotype $u'$ given a particular midparent phenotype $z$:

$$u'(z) = h_i^2(z-u_i) + u_i, \tag{S36}$$

where $u_i$ is the mean phenotype of the population of species $i$, and $h_i^2$ is the heritability given as $\frac{V_i}{\rho_i}$, i.e., ratio of genetic variance by phenotypic variance. The variance of the offspring distribution, which is also called the segregation variance or residual variance (Lande 1981, Slatkin & Lande 1994, Roughgarden 1972, Taper & Chase 1985), can be written as:

$$\rho_s = (1-\frac{h_i^2}{2})V_i + \epsilon = \rho_i - \frac{h_i^2}{2}V_i. \tag{S37}$$

Here, $\epsilon$ is the environmental variance, and $\rho_i$ is the phenotypic variance.

The above expression can be derived as follows. The total variance of a dependent variable, i.e., the variance of all offspring is equal to the explained variance plus the residual variance. The residual variance is also known as the segregation variance, which here we write as $\rho_s$.

Reanalysing equation S36 and dropping the subscript $i$ for convenience (for now) we can rewrite that as:

$$u'(z) = h^2 z + (1-h^2)u, \tag{S38}$$

which is a simple linear regression of the form $y = az + b$. Here, $a = h^2$, and $b = (1-h^2)u$, and $y = u'(z)$. For this ordinary least squares regression line, $b$ can be estimated as $b = \bar{y} - a\bar{z}$, $\bar{y}$ is the mean for variable $y$, and $\bar{z}$ is the mean of variable $z$. This is equivalent to for equation S38:

$$(1-h^2)u = \bar{u} - h^2\bar{z}, \tag{S39}$$

where $\bar{u}$ is the mean of expected phenotype offspring distribution. Now from standard linear regression analysis, $a = h^2 = \frac{S_{uz}}{S_{zz}}$, where, $S_{uz}$ is the covariance of midparent phenotype and offspring phenotype, and $S_{zz}$ is the variance of the midparent phenotype. Thus, $S_{uz} = \frac{1}{2}V$, and $S_{zz} = \frac{1}{2}\rho$. Next, we then arrive at the segregation variance or the residual variance (which is also known as residual sum of squares, RSS), and can be written as:

$$\rho_s = \sum_j^n \big(u_j - u'(z_j)\big)^2 = \sum_j^n \big(u_j - h^2(z_j - u) - u\big)^2 = \sum_j^n \big(u_j - h^2 z_j - u(1-h^2)\big)^2. \tag{S40}$$

Here, $u_j$ is the jth value of the variable to be predicted, $z_j$ is the jth value of the explanatory variable, and $u'(z_j)$ is the expected or predicted value of $u_j$. Rearranging equation S40 and substituting equation S39 we get

$$\sum_j^n \big(u_j - h^2 z_j - u(1-h^2)\big)^2 = \sum_j^n \big(h^2(\bar{z} - z_j) - (\bar{u} - u_j)\big)^2 = \tag{S41}$$

$$\sum_j^n \Big((h^2)^2(\bar{z} - z_j)(\bar{z} - z_j) - 2h^2(\bar{z} - z_j)(\bar{u} - u_j) + (\bar{u} - u_j)(\bar{u} - u_j)\Big). \tag{S42}$$

The above equation S41 can be further simplified after substituting $h^2 = \frac{S_{uz}}{S_{zz}}$ to:

$$\rho_s = (h^2)^2 S_{zz} - 2h^2 S_{zu} + S_{uu} = -h^2 S_{zu} + S_{uu} = S_{uu}\Big(1 - \frac{S_{zu}^2}{S_{zz}S_{uu}}\Big) \tag{S43}$$

With this, we finally arrive at the segregation variance denoted in equation S37:

$$\rho_s = S_{uu}\Big(1 - \frac{S_{zu}^2}{S_{zz}S_{uu}}\Big) = \rho\Big(1 - \frac{h^2}{2}\frac{V}{\rho}\Big) = \rho - \frac{h^2}{2}V \tag{S44}$$

since, $\frac{S_{zu}}{S_{zz}} = h^2$, and since $S_{uu}$ is just the offspring phenotypic variance which amounts to $\rho$, then $\frac{S_{zu}}{S_{uu}} = \frac{1}{2}\frac{V}{\rho}$, as $S_{zu} = \frac{V}{2}$.

Since, $\rho_i$ is the phenotypic variance which is given as (here we revert back to using subscript $i$ again which denotes for a species $i$ in a mutualistic community):

$$\rho_i = V_i + \epsilon. \tag{S45}$$

The total phenotypic variance of all the offspring, say $\rho_i'$, will be the sum of the segregation variance and the variance of the expected phenotype given the mid-parent phenotype (Taper & Chase 1985) which can be written as

$$\rho_i' = Var[u_i'(z)] + \rho_s^2 = Var[h_i^2(z - u_i) + u_i] + \rho_s^2 \tag{S46}$$

$$= h_i^4 Var[z] + \rho_i - \frac{h_i^2}{2}V_i \tag{S47}$$

Here we used the fact that $Var(a + bX) = b^2 Var(X)$. Substituting equation S45 in the above equation we get,

$$V_i' + \epsilon = h_i^4 Var[z] + V_i + \epsilon - \frac{h_i^2}{2}V_i \tag{S48}$$

$$V_i' - V_i = h_i^4 Var[z] - \frac{h_i^2}{2}V_i \tag{S49}$$

This equation is similar to change in phenotypic variance without any dominance or linkage disequilibrium as derived in Walsh & Lynch (2018)(pg 552-554). Finally, we if write $\Delta V = V' - V$ as the difference in variance in two generations, we can write the above equation as:

$$\Delta V = h_i^2\left(h_i^2 Var[z] - \frac{1}{2}V_i\right). \tag{S50}$$

Note that the above equation of genetic variance change is valid but needs to be adjusted for the variance part. We need to consider what would happen to the variance of parental phenotype after selection. If in the population mating is random then we can explicitly write the variance of the mid-parent phenotype from Taper & Chase (1985) formulation as

$$Var[z] = \frac{1}{2}\widetilde{\rho}, \tag{S51}$$

where $\widetilde{\rho}$ is the phenotypic variance after selection. Thus, we can re-write equation S50 by substituting equation S51 as:

$$\Delta V_i = \frac{h_i^2}{2}\left(h_i^2\widetilde{\rho} - V_i\right). \tag{S52}$$

Thus we can finally arrive at the equation of phenotypic variance dynamics by substituting equation S34 into equation S52 (Taper & Chase 1985, Barabás *et al.* 2022):

$$\Delta V_i = h_i^2 \frac{1}{2}\left[ h_i^2 \left( \rho_i + s \int \left( (z-u_i)^2 - \rho_i \right) r_i(z) p_i(z) dz \right) - V_i) \right] + O(s^2), \tag{S53}$$

where index $i$ is for species $i$.

Now substituting, $V_i = h_i^2 \rho_i$ into equation S53 we get:

$$\Delta V_i = s \cdot \frac{1}{2} \left( \int h_i^4 \left[ (z-u_i)^2 - \rho_i \right] r_i(z) p_i(z) dz \right) + O(s^2). \tag{S54}$$

Similar to Bürger (2011), pg 3-6, we now will arrive at the dynamics of genetic variance. Selection intensity $s$ has units of generation time. We let the generation time $s$ go to zero to finally arrive at the dynamics of variance. In the weak selection limit, since $r_i(z,t)$ in equation S34 is the per-capita growth rate with units of inverse of generation (if time is measured in generation), then selection intensity $s$ has units of generation. The change in genetic variance of the phenotype is approximately proportional to the product of the selection intensity $s$ and a functioin of per-capita growth rate (see equation S54). Now, suppose $s$ is small, for example 0.01, then per-generation change in variance of phenotype is also very small i.e., 1%. Only when a substantial number of generations has passed, we would observe a significant change in the mean and variance of the phenotype. Thus in the limit

$$\frac{dV_i}{dt} = \lim_{s \to 0} \frac{V_i(t+s) - V_i(t)}{s} = \lim_{s \to 0} \frac{\Delta V_i}{s} = \frac{1}{2} \left( \int h_i^4(t) \left[ (z-u_i)^2 - \rho_i \right] r_i(z,t) p_i(z,t) dz \right). \tag{S55}$$

Solving the above integrals, we can thus write the changes in genetic variance over time as:

$$\frac{dV_i}{dt} = \frac{1}{2} V_i^2 \left( v_i(T) - \sum_{i \neq j}^{S_A} \Gamma_{ij}^A N_j^A + \nu_i \right), \tag{S56}$$

where we have used that $h_i^4 = (\frac{V_i}{\rho_i})^2$ and

$$v_i(T) = \int \frac{1}{\rho_i^2} \left( (z-u_i)^2 - \rho_i \right) b(z,t) p_i(z) dz = \frac{2g_i \left( 2u_i^2 - 4Tu_i + 2T^2 - 2b_w^2 - \rho_i \right)}{\sqrt{(b_w^2 + \rho_i)}} \frac{1}{(2b_w^2 + \rho_i)} \exp\left(-\frac{(T-u_i)^2}{(2b_w^2 + \rho_i)}\right) \tag{S57}$$

is the evolutionary impact on changes in genetic variance due to tolerance to temperature,

$$\Gamma_{ij}^A = \int \int \frac{1}{\rho_i^2} \left( (z-u_i)^2 - \rho_i \right) \alpha(z, z''') p_i(z) p_j(z''') dz dz''' = \frac{\alpha_{ij}^A}{(\rho_i + \rho_j + \omega_c^2)^2} \left( (u_i - u_j)^2 - (\rho_i + \rho_j + \omega_c^2) \right) \tag{S58}$$

is the evolutionary impact on genetic variance due to interspecific competition on the same guild of species, and finally

$$\nu_i = \int \int \frac{1}{\rho_i^2} \left( (z-u_i)^2 - \rho_i \frac{\sum_k \gamma(z,z') A_{ik} N_k^{(P)}(t)}{1 + H \int \sum_k \gamma(z,z'') A_{ik} N_k^{(P)}(t) p_k^P(z'',t) dz''} p_k^P(z',t) dz' \right) p_i^A(z,t) dz \tag{S59}$$

is the evolutionary impact due to mutualistic interaction. $\nu_i$ is analytically not solvable and hence we resort to solving it numerically.

### 3 Supplementary Figures:

### References


Barabas, G. & D'Andrea, R. (2016) The effect of intraspecific variation and heritability on community pattern and robustness. *Ecology Letters* **19**, 977–986.

Barabás, G., Parent, C., Kraemer, A., Van de Perre, F. & De Laender, F. (2022) The evolution of trait variance creates a tension between species diversity and functional diversity. *Nature Communications* **13**, 2521, number: 1 Publisher: Nature Publishing Group.

Baruah, G. (2022) The impact of individual variation on abrupt collapses in mutualistic networks. *Ecology Letters* **25**, 26–37, _eprint: https://onlinelibrary.wiley.com/doi/pdf/10.1111/ele.13895.

Baruah, G. & Wittmann, M. (2024) Reviving collapsed plant–pollinator networks from a single species. *PLOS Biology* **22**, e3002826, publisher: Public Library of Science.

Bürger, R. (2011) *Some Mathematical Models in Evolutionary Genetics*, pp. 67–89. Springer Basel, Basel.

Falconer, D.S. & Mackay, T.F.C. (1996) Introduction to quantitative genetics. *Introduction to quantitative genetics* .

Lande, R. (1976) Natural selection and random genetic drift in phenotypic evolution. *Evolution* pp. 314–334.

Lande, R. (1981) THE MINIMUM NUMBER OF GENES CONTRIBUTING TO QUANTITATIVE VARIATION BETWEEN AND WITHIN POPULATIONS. *Genetics* **99**, 541–553.

Nagylaki, T. (1993) The evolution of multilocus systems under weak selection. *Genetics* **134**, 627–647.

Roughgarden, J. (1972) Evolution of Niche Width. *The American Naturalist* **106**, 683–718, publisher: The University of Chicago Press.

Slatkin, M. & Lande, R. (1994) Segregation variance after hybridization of isolated populations. *Genetics Research* **64**, 51–56.

Taper, M.L. & Chase, T.J. (1985) Quantitative Genetic Models for the Coevolution of Character Displacement. *Ecology* **66**, 355–371, _eprint: https://onlinelibrary.wiley.com/doi/pdf/10.2307/1940385.

Walsh, B. & Lynch, M. (2018) *Evolution and selection of quantitative traits.* Oxford University Press.

Wild, G. & Traulsen, A. (2007) The different limits of weak selection and the evolutionary dynamics of finite populations. *Journal of Theoretical Biology* **247**, 382–390.

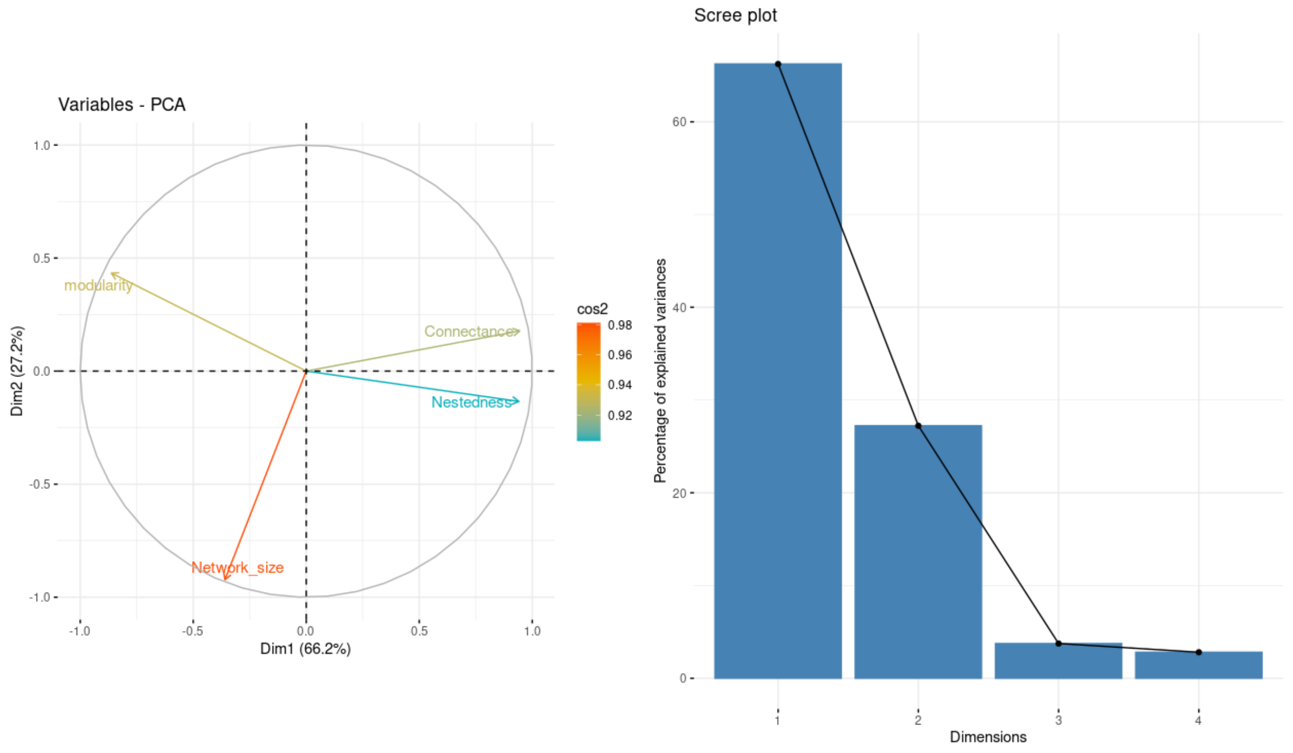


Figure S1: Principal component analysis of four network metrics: nestedness (NODF), network size, connectance and modularity. (Left) PC1 is correlated positively to connectance and nestedness and negatively to modularity, and PC2 is negatively correlated to network size. (Right) PC1 and PC2 explain 99 % of variation.

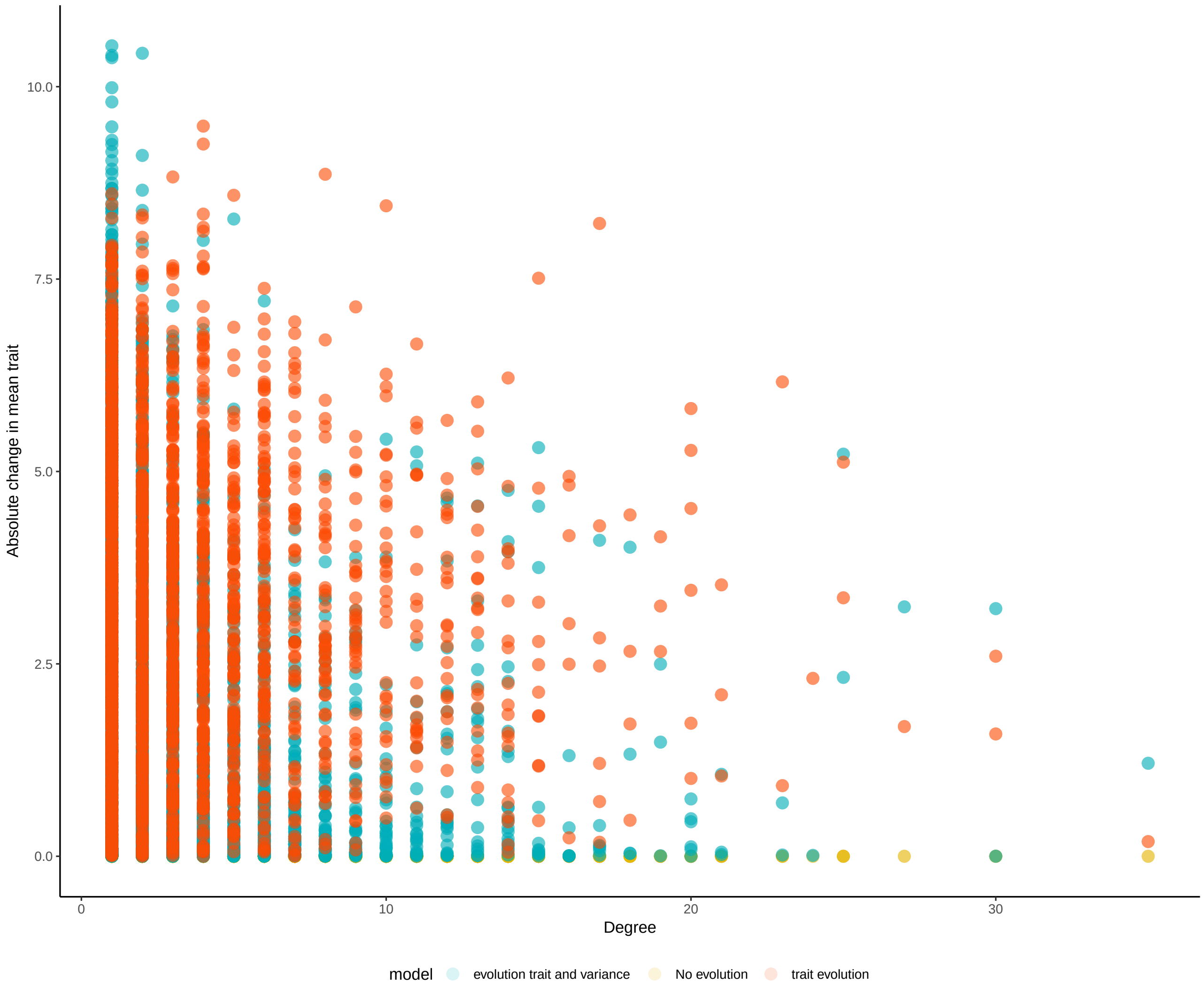


Figure S2: Absolute change in species mean trait in relation to species degree for the three different models. With evolution of mean trait and genetic variance, changes in mean trait ranged to as high as +10, while changes were smaller when only the mean trait evolved in response to abrupt shift in temperature.

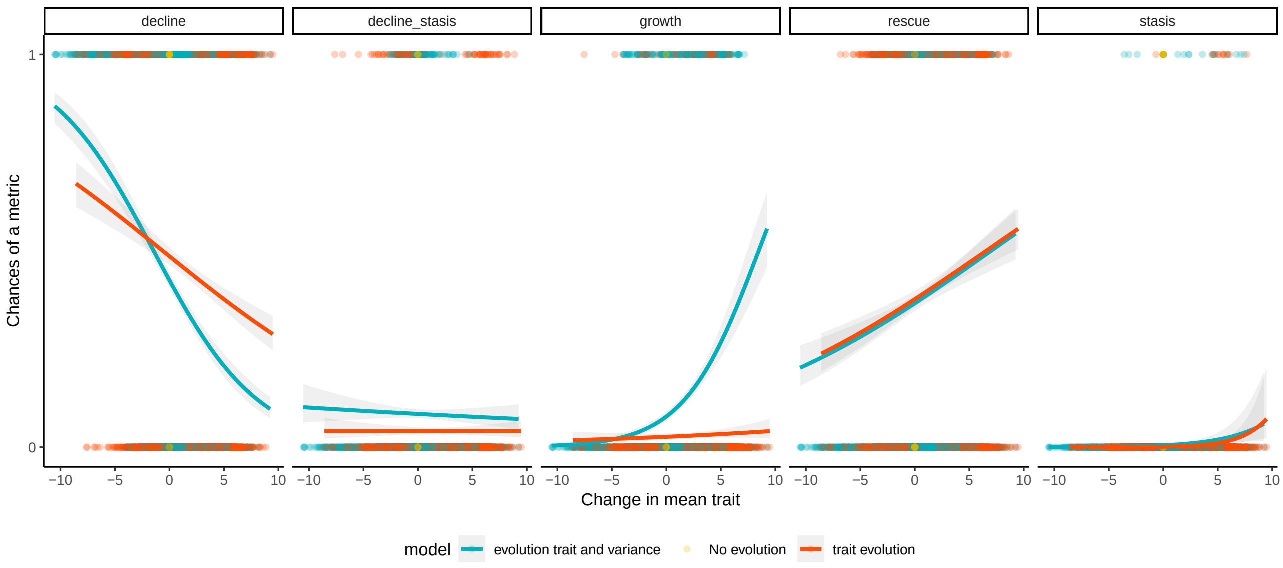


Figure S3: Chances of species decline, decline-stasis, growth, rescue, and stasis for the three different models in relation to change in mean trait. Higher positive change in mean trait was related to a smaller probability of decline, high chances of rescue and high chances of growth.

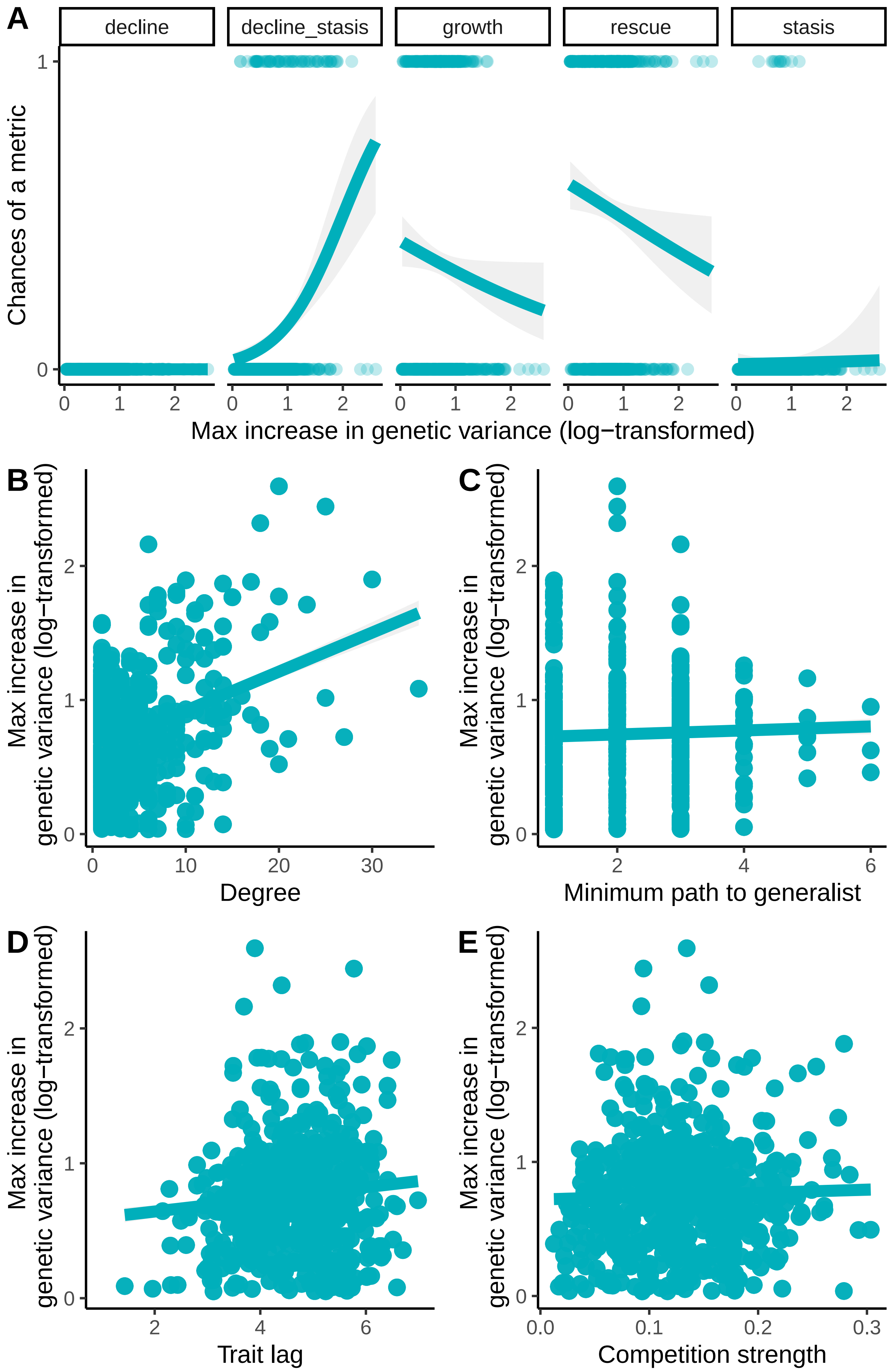


Figure S4: Chances of the different outcomes and maximum increase in genetic variance for the model when both mean and genetic variance of species evolve in response to abrupt change in temperature. A) Chances of decline-stasis increases as maximum increases in genetic variance is reached for species. In addition, chances of evolutionary rescue also increase as species exhibit larger increases in genetic variance. B) However, maximum increase in genetic variation exhibited by species remains positively correlated with species degree, and C) slightly positively correlated to the distance to a hyper-generalist species. Maximum increases in genetic variance for species reached was positively correlated with (D) species initial trait lag and (E) competition faced by a species.

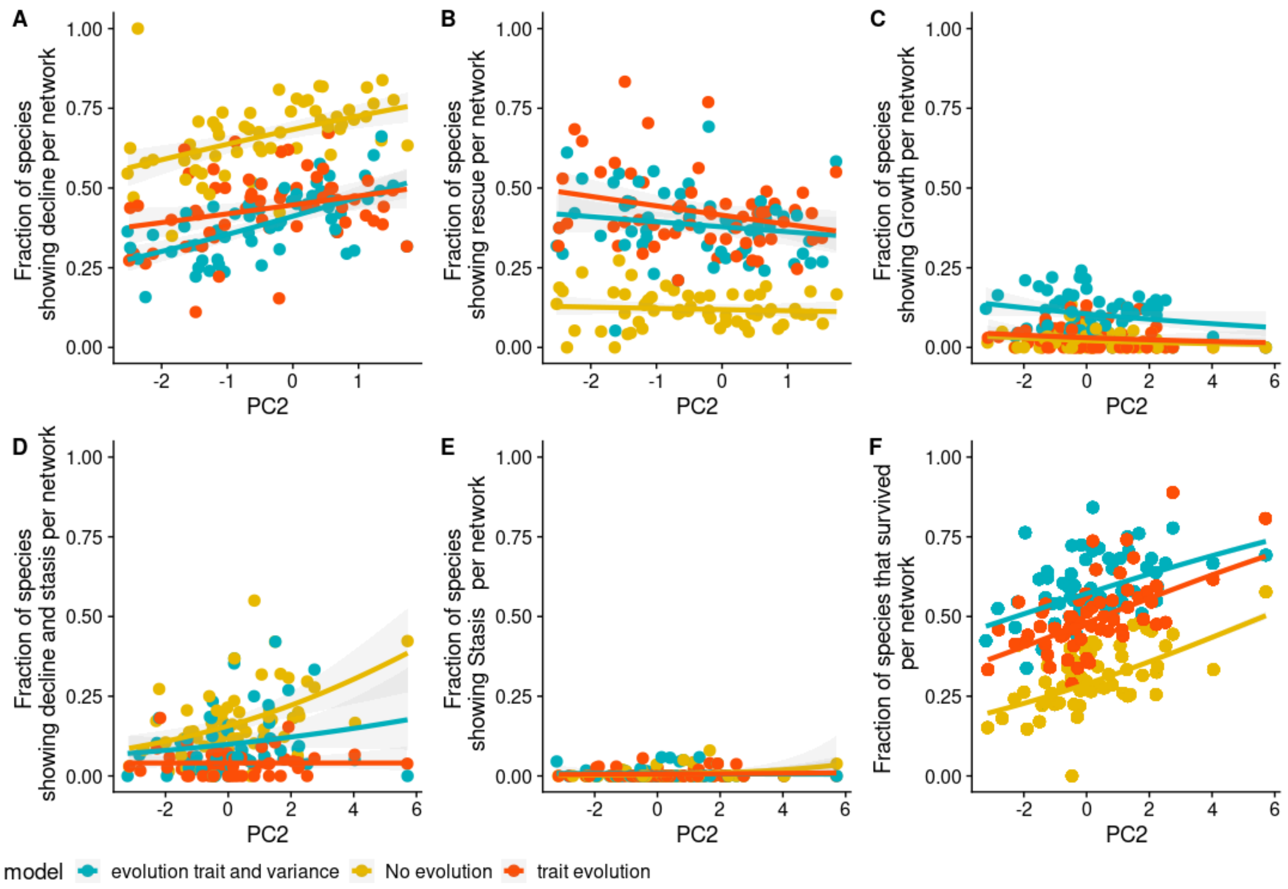


Figure S5: Relationship between proportion of species showing (A) decline, (B) rescue, (C) growth, (D) decline-stasis, (E) stasis, (F) survival in relation to PC2 which correlates negatively with network size i.e, low PC2 values correlates with large networks. Parameters as in table 1.

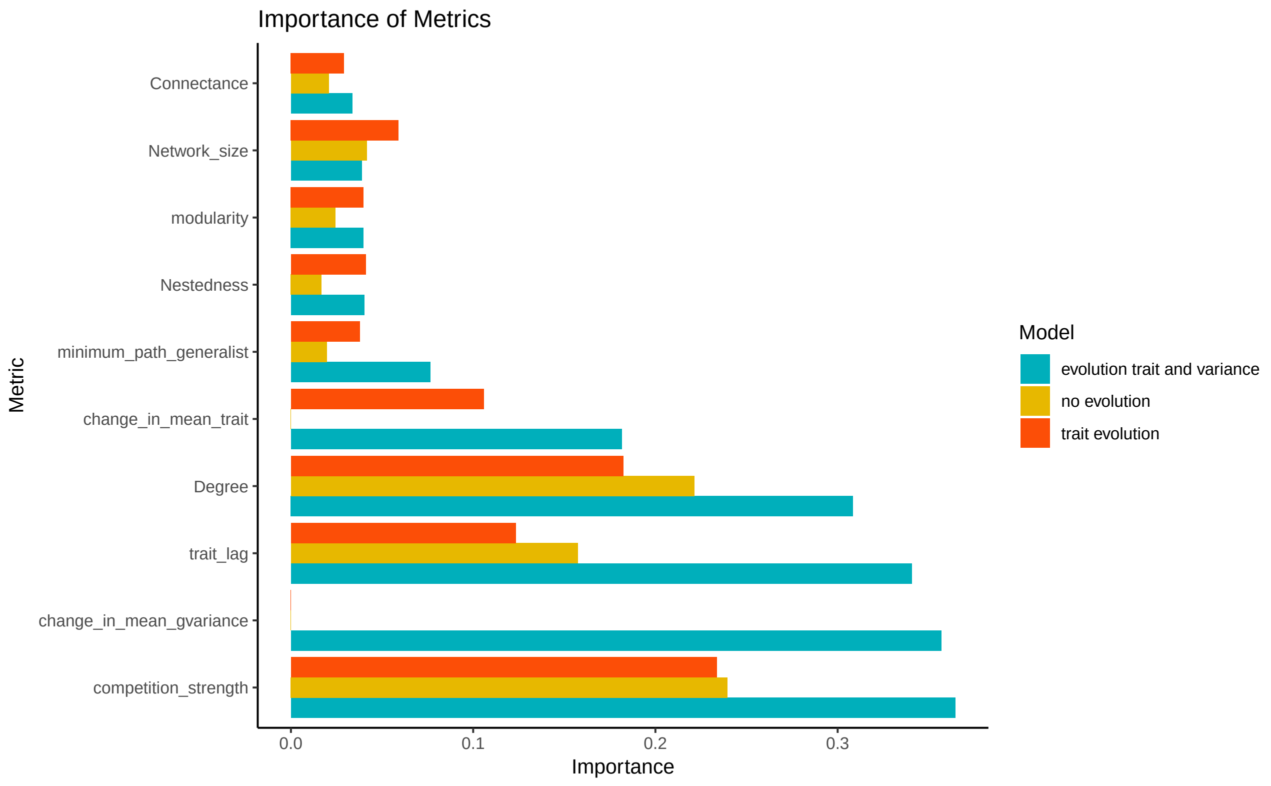


Figure S6: Importance of all the metrics analysed for our multivariate binomial random forest models in relation to the three models analysed. Parameters as in table 1. Competition strength, trait-lag, species degree and minimum path to a super generalist are important in multivariate response of species to abrupt change in environmental temperature.

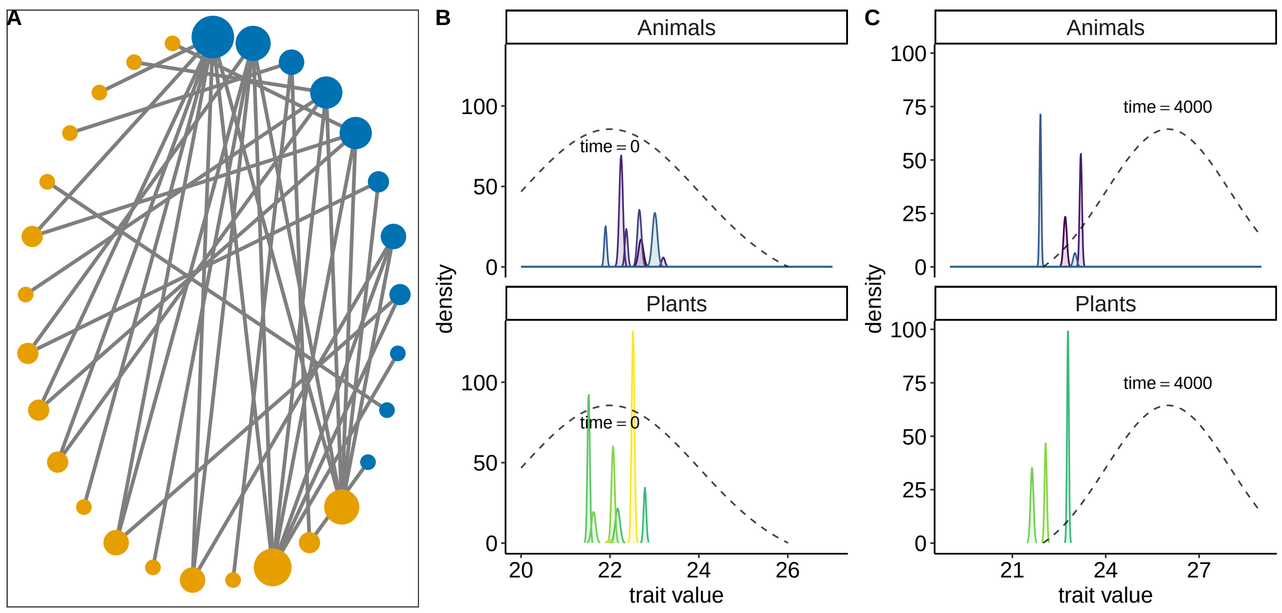


Figure S7: Response of species after a shift in the environmental optima from 22°C to 26°C when there is no evolutionary dynamics at play either in the mean phenotype or in species' genetic variance. A) 29 species plant-pollinator network. B) At eco-evolutionary equilibrium, species arrange themselves in the trait axis, with the dashed curve line representing the temperature tolerance curve. C) After the shift to a new temperature optima of 26° C, some species exhibit the behavior of growth, decline-stasis, and one species show rescue behaviour. The growth behaviour was shown by one species which had low trait-lag, and after the shift to the new optimum, it gained fitness. A lot of the species exhibited decline-stasis behavior, and only one species exhibited rescue behavior. Parameters as in table 1.

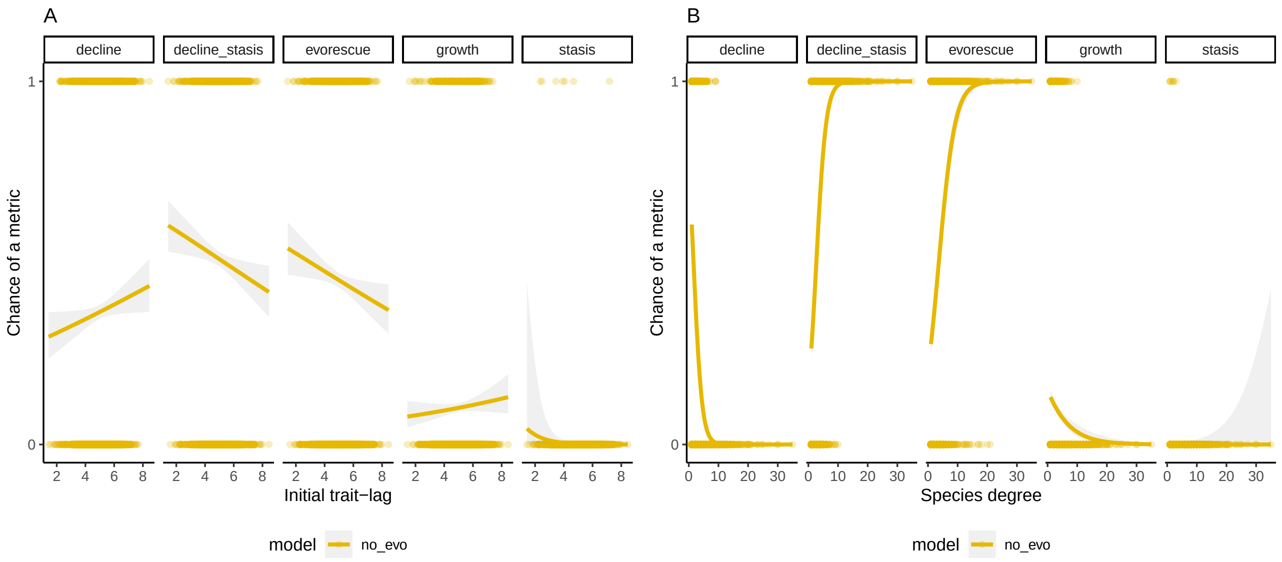


Figure S8: Chance of decline, decline-stasis, rescue, growth, and stasis in relation to A) species trait lag, (B) species degree, when species do not evolve in either the mean trait or in genetic variance. Lines represent predicted response of the different metrics of eco-evolutionary response of species as given by a generalised linear model with binomial error distribution, and shaded regions indicate 95% confidence interval. Parameters as in Table 1.

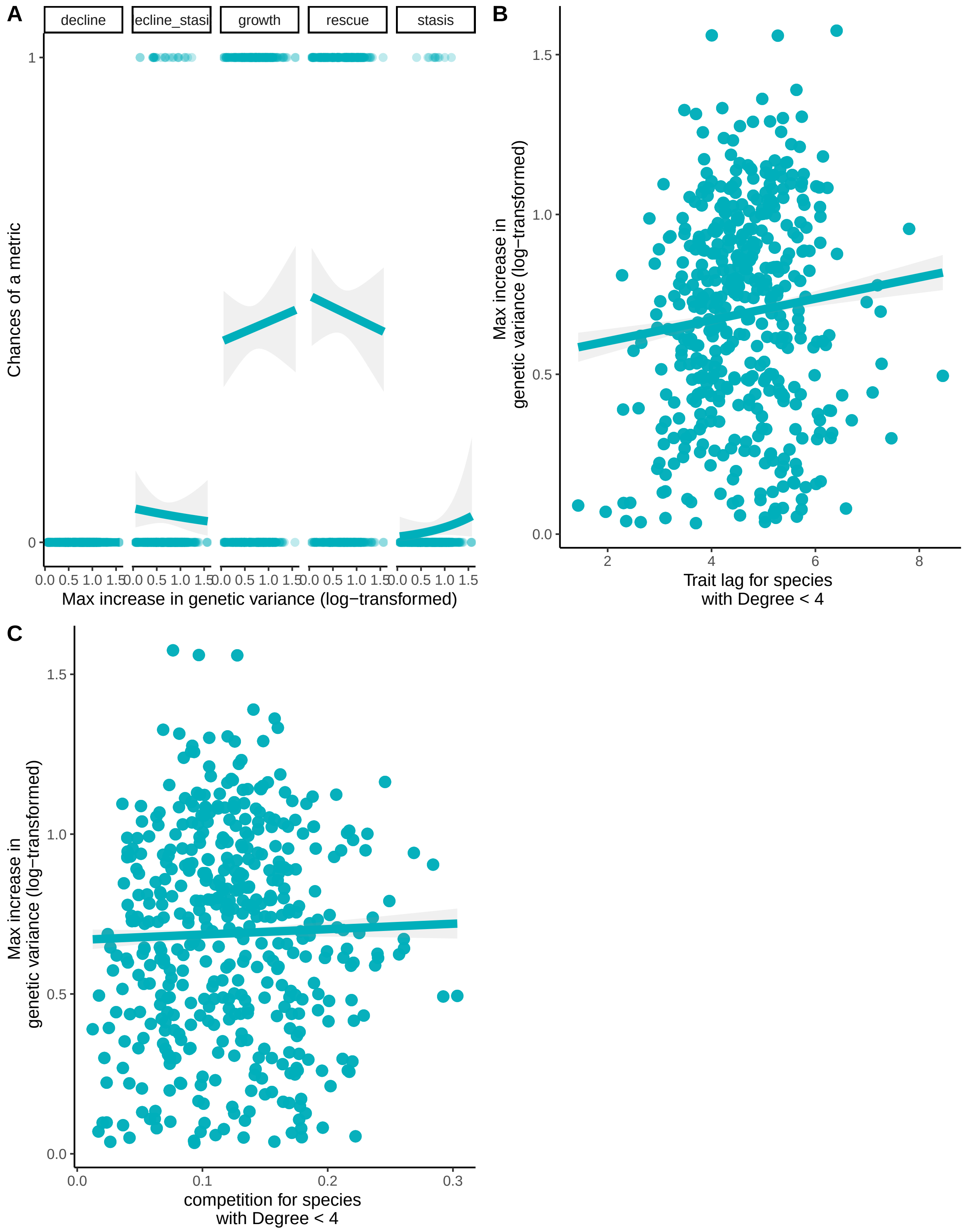


Figure S9: A) Chance of decline, decline-stasis, growth, rescue, and stasis in relation to maximum increases of genetic variance for specialist species, i.e., degree $< 4$. (B) Increases in genetic variance are directly related to trait-lag, and slightly positively related to C) competition they faced. In (A) lines represent predicted response of the different metrics of eco-evolutionary response of species as given by a generalised linear model with binomial error distribution, and shaded regions indicate 95% confidence interval. In (B-C) lines represent linear regression with 95 % confidence interval. Parameters as in Table 1.

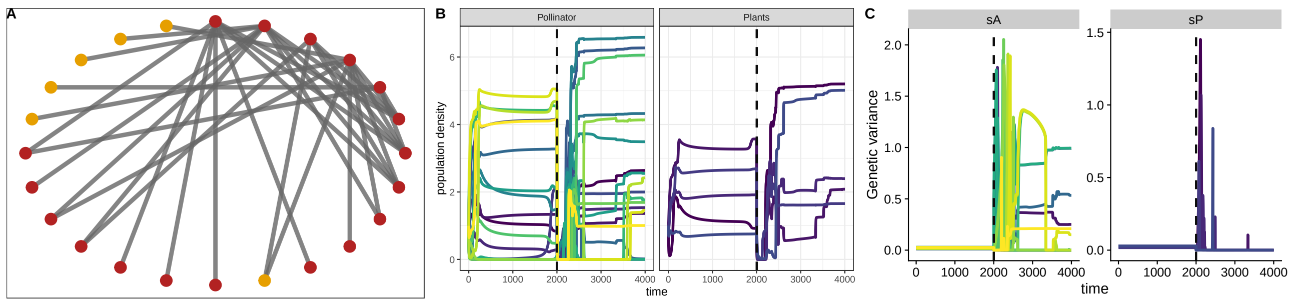


Figure S10: Dynamics of genetic variance and population dynamics of an example 23 species plant-pollinator network after an environmental temperature shift. A) 23 species plant-pollinator network. Red nodes indicate the species that survived after a temperature shift. B) Population dynamics of plant-pollinator network. C) Evolutionary dynamics of genetic variance of 23 species plant-pollinator networks. After a shift in the temperature, genetic variance increases rapidly which aids in survival of species. Parameters as in Table 1.

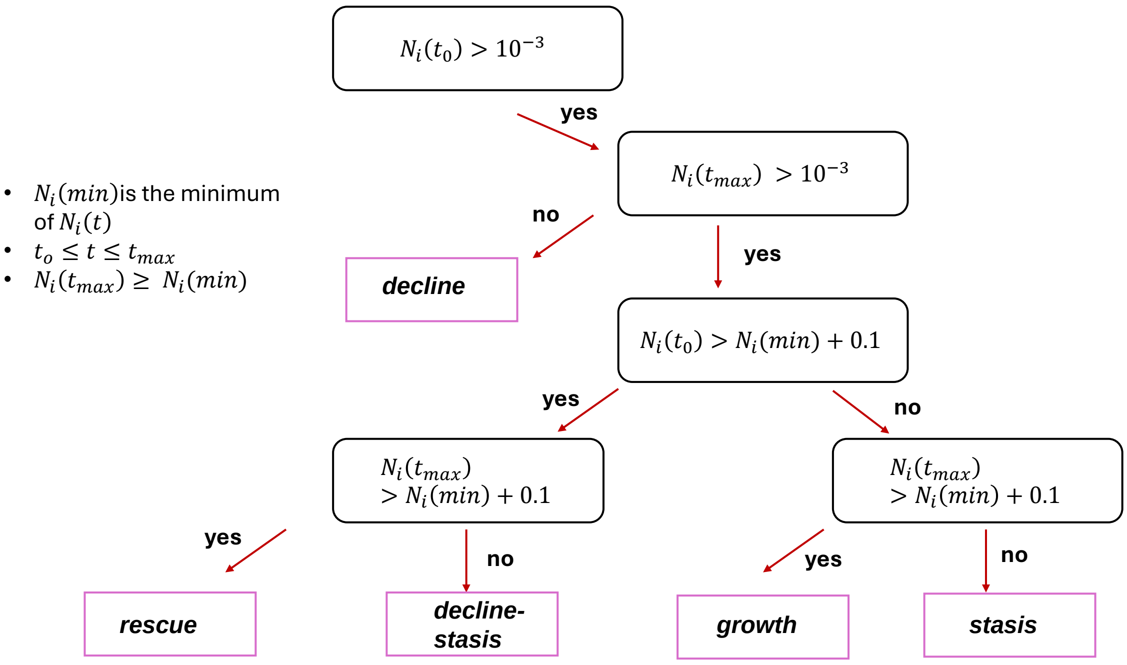


Figure S11: Classification used to determine from simulations which species in a network can be categorized in either decline, rescue, decline-stasis, growth or just stasis. This figure is in regards to figure 1 in the main text. $t_0$ is the time point at which the local environmental temperature shifts, and $t_{max}$ is the final time point of our simulations. $N_i(min)$ is the minimum population size of species $i$ in the time interval $10 < t < t_{max}$.

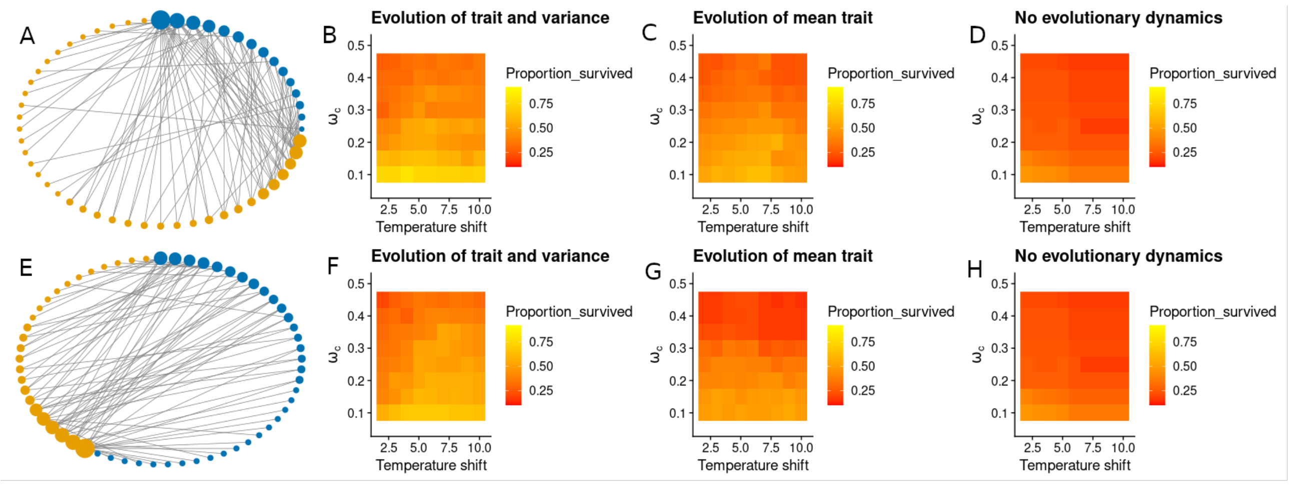


Figure S12: Parameter space for strength of species competition $\omega_c$ and the magnitude of temperature shift on final survival of species, for three models and two contrasting plant-pollinator networks. (B,F) When mean trait and genetic variance evolve, the proportion of species that survive is higher in comparison to the other two models (C,G) evolution of only mean trait, and (D,H) no evolution, across the two different networks. Even during an unrealistically large temperature shift of 10° C at low competition strength $\omega_c = 0.1$, we observe almost 60% survival of species when both mean trait and variance evolves. Parameters as in table 1.
